# PIE: Generalizing perturbation effects across unseen perturbations, contexts and datasets

**DOI:** 10.64898/2026.10.02.756297

**Authors:** Rishi Verma, Abhinav Adduri, Beatrice Bevilacqua, Basak Eraslan, Dave P. Burke, Hani Goodarzi, Yusuf H Roohani

**Author notes:** Corresponding authors: Y.H.R.; H.G. These authors supervised this work (Dave P. Burke, Hani Goodarzi, Yusuf H Roohani).

## Abstract

Predicting cellular response to perturbations is key to understanding biological mechanisms and selecting therapeutic targets. However, generalizing across cellular contexts, perturbations, and experimental datasets remains challenging due to incomplete representations of the system being perturbed and the difficulty of measuring true perturbation effects within that system. We introduce PIE, a model that addresses these limitations by reformulating the learning task around population-level perturbation effects and incorporating auxiliary inputs that better characterize the biological system. PIE directly predicts differentially expressed genes and changes in gene expression for each (context, perturbation) pair, using external biological knowledge, baseline gene expression, and observed perturbation responses. Its architecture accommodates input sources with variable feature sets and predicts effects on genes seen and unseen during training. On the Replogle–Nadig dataset, PIE achieves state-of-the-art performance on most metrics across all tested generalization settings, including the most challenging setting where both the context and perturbation are unseen during training. For predicting differentially expressed genes, PIE achieves 1.2–3.2 times the AUPRC of the strongest baseline in each setting: across unseen contexts, perturbations, experimental datasets, and jointly unseen contexts and perturbations. Overall, PIE provides a framework for generalizing perturbation effects across diverse experimental conditions.

## 1. Introduction

Understanding how cells respond to genetic and chemical perturbations is important for uncovering the mechanisms that drive biological systems and disease. Every perturbation experiment is defined by the cellular context in which the perturbation is performed, the applied perturbation itself, and the broader experimental context or dataset. Computational models can help guide experiments by predicting responses under conditions not already tested. However, generalizing across variations in these experimental conditions requires distinguishing the biological effects of the perturbation from experiment-specific factors. This remains challenging for two related reasons: incomplete representations of the biological system being perturbed and the difficulty of measuring the true effects of perturbation within that system. Most existing perturbation prediction models rely mainly on transcriptomic data or a few auxiliary data sources to represent contexts or perturbations. Moreover, they attempt to predict perturbation effects at the level of individual cells, which is challenging because current single-cell perturbation technologies cannot directly measure perturbation effects at this level. This mismatch is also reflected in how these models are evaluated, as most evaluation metrics focus on population-level measurements, such as the set of differentially expressed genes following perturbation (Adduri et al., 2026).

We introduce PIE (Perturbation is Everything), a model that addresses these challenges by simplifying the learning task and incorporating auxiliary inputs that better characterize the biological system. Unlike some existing perturbation prediction models that attempt to predict the absolute gene expression state of a cell following perturbation, PIE focuses on learning population-level effects for each condition. This provides a more direct training signal for perturbation effects and facilitates generalization to new conditions. Thus, the input is a (context, perturbation) pair and the task is to predict condition-level outcomes such as the set of differentially expressed genes, the direction of differential expression, and the binned magnitude of mean expression change. This contrasts with predicting individual cell-level responses that are harder to measure precisely and are less consistent across experiments. By focusing on condition-level effects, PIE greatly reduces the space of possible outcomes that the model must predict compared to conventional transcriptomic models, thereby enhancing transferability to new conditions.

To build a more complete representation of the system being perturbed, PIE leverages three kinds of auxiliary inputs: knowledge sources, which embed each context and perturbation using external databases such as Cellosaurus, NCBI Gene, and PubChem (Bairoch (2018),Brown et al. (2015),Kim et al. (2025)); the mean control expression of the query context; and summary statistics of gene-level responses from experimental conditions observed during training. To accommodate these heterogeneous inputs across datasets and perturbation modalities, PIE builds on Perceiver IO (Jaegle et al., 2022). Source-specific encoders let PIE combine inputs whose feature sets vary in number, type, and dimensionality. This enables a common architecture to operate across diverse datasets such as chemical and genetic perturbation experiments. Because each output gene is represented using its description and available experimental evidence, PIE can predict outcomes for both seen and unseen genes during training. This allows PIE to accommodate multiple datasets without normalizing gene vocabularies.

Unlike existing baselines, PIE can generalize across multiple axes of experimental covariates: dataset, context, and perturbation, as well as combinations of these axes. Across each axis, PIE outperforms or matches existing baselines that are specialized for that axis. We observe the largest gains in AUPRC for classifying differentially expressed genes, where PIE outperforms the strongest evaluated baseline in each setting by 1.2–3.2 times across within- and cross-dataset generalization tasks. PIE’s modular architecture further enables ablation studies that assess the relative contributions of different input sources to model performance across generalization settings and evaluation metrics.

## 2. Related Work

Predicting transcriptional responses to interventions encompasses several distinct generalization problems. Methods for context generalization transfer the effect of a known perturbation to a new background, either by transporting its unperturbed cells (Bunne et al., 2023; Klein et al., 2025; Adduri et al., 2026; Dong et al., 2026) or by learning how population-level responses vary across contexts (Svensson et al., 2026; Palla et al., 2026). Methods for perturbation generalization instead replace learned perturbation identities with representations derived from prior knowledge, such as molecular structure for compounds (Hetzel et al., 2022) and gene interaction graphs, text descriptions, or pretrained gene embeddings for genetic perturbations (Roohani et al., 2024; Chen and Zou, 2025; Littman et al., 2025; Wenkel et al., 2026; Liang and Singh, 2026). Transfer across datasets remains difficult even for models trained on hundreds of millions of cells (Adduri et al., 2026).

Methods also differ in the resolution at which they model perturbation effects. Single-cell foundation models such as scGPT (Cui et al., 2024), Geneformer (Theodoris et al., 2023), scFoundation (Hao et al., 2024) and UCE (Rosen et al., 2026) are pretrained on large observational atlases to learn general-purpose representations of transcriptomic state, and are fine-tuned or prompted to predict perturbed expression profiles. Generative and transport-based models learn the distribution of perturbed cells directly (Lotfollahi et al., 2023; Bunne et al., 2023; Klein et al., 2025; Palla et al., 2025). Because the unperturbed state of a perturbed cell is never observed, these models must separate the perturbation effect from heterogeneity within the cell population and from technical variation across experiments, which motivates set-based architectures (Adduri et al., 2026) and disentangled or sparse latent representations (Bereket and Karaletsos, 2023; Piran et al., 2024). Others such as PRESAGE (Littman et al., 2025), COMPASS (Liang and Singh, 2026) and Rhaister-O (Svensson et al., 2026) treat the condition label (context, perturbation) as the sample and predict pseudobulk responses.

## 3. Method

### 3.1. Problem setup

A context *c* is the cellular background in which a perturbation is applied. In our datasets, a context is a cell line, either unstimulated, or treated with a signaling molecule such as a cytokine. A perturbation *p* is a CRISPRi knockdown of a target gene or a small molecule at a given dose. An experimental condition is a pair (*c, p*) measured in a dataset, and we write *C*_*cp*_ for its set of profiled cells. Each dataset also includes unperturbed control cells *C*_*c*,_ctrl for every *c*. Different datasets may measure different sets of genes, and we use *G* to denote the size of the measured gene vocabulary.

For every experimental condition and measured gene *g*, we derive three labels by comparing *C*_*cp*_ with *C*_*c*,_ctrl (Appendix A.1). Let *n*_*ig*_ be the normalized expression of gene *g* in cell *i, x*_*ig*_ = log (1 +*n*_*ig*_) its lognormalized expression, and 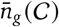 and 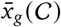 their means over a set of cells *C*. The DE label *y*_*cpg*_ ∈ {0, 1} indicates whether a Wilcoxon rank-sum test yields a Benjamini–Hochberg FDR below 0.05. The LFC and the pseudobulk delta expression are:

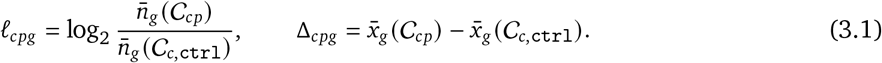

The LFC is defined only for DE genes, and its sign gives the direction of change. All three labels are derived from cell populations rather than individual cells.

During training, the labels are observed for a training set *T* of experimental conditions. At test time, the model must predict *y*_*cpg*_, ℓ_*cpg*_, and Δ_*cpg*_ for every measured gene of a condition (*c, p*) ∉ *T*, using only information that is available without measuring that condition, such as external knowledge about *c* and *p*, the control cells of *C*_*c*, ctrl_, and the responses in *T*. The generalization regime depends on what *T* contains. A context is unseen if no condition in *T* involves *c*, and a perturbation is unseen if no condition in *T* involves *p*. The two may also be unseen jointly. In all regimes, model parameters stay fixed at test time.

### 3.2. Knowledge Sources

To make predictions for contexts and perturbations that have never been measured, PIE represents each context and perturbation using information from external databases, referred to as Knowledge Sources. A Knowledge Source maps a context or perturbation to a single pooled embedding or a sequence of embeddings (Appendix A.2). We project each embedding to the model width, add a learned embedding that identifies the type of knowledge source, and concatenate the embeddings from all sources of an experimental condition into a single sequence of input embeddings. When a database has no entry for a context or perturbation, that source is omitted from the sequence (Appendix A.2.2).

Contexts are represented by a single source, the Context Description, which is the cell line’s Cellosaurus entry (Bairoch, 2018), embedded with a text embedding model. For the stimulated contexts of Jiang et al., we append a description of the stimulus to the entry.

Perturbations are represented by a richer set of Perturbation Knowledge Sources. Every perturbation has a Perturbation Description, embedded in the same way as the Context Description: the NCBI Gene entry of the target gene for genetic perturbations (Brown et al., 2015), and the PubChem entry of the compound, together with its dose, for chemical perturbations (Kim et al., 2025). For genetic perturbations, we add four sources that describe the target gene: NCBI Gene Annotations, which list its Gene Ontology terms (Ashburner et al., 2000; The Gene Ontology Consortium et al., 2025), disease associations (McKusick-Nathans Institute of Genetic Medicine, Johns Hopkins University, 2026), and gene-group memberships; ESM2 embeddings of its protein sequence (Lin et al., 2023); STRING embeddings of its protein interaction network (Szklarczyk et al., 2025; Hu et al., 2025); and DepMap profiles of its knockout effect on viability across cell lines (DepMap, Broad, 2025; Dempster et al., 2021). For chemical perturbations, we add ChemBERTa embeddings of the compound’s SMILES string (Ahmad et al., 2022) and three measured activity profiles: transcriptional responses from L1000 (Subramanian et al., 2017), viability effects from PRISM (Corsello et al., 2020), and morphological effects from JUMP (Chandrasekaran et al., 2023). Coverage varies across sources (Appendix Table 1), particularly for the measured compound profiles.

**Table 1.**
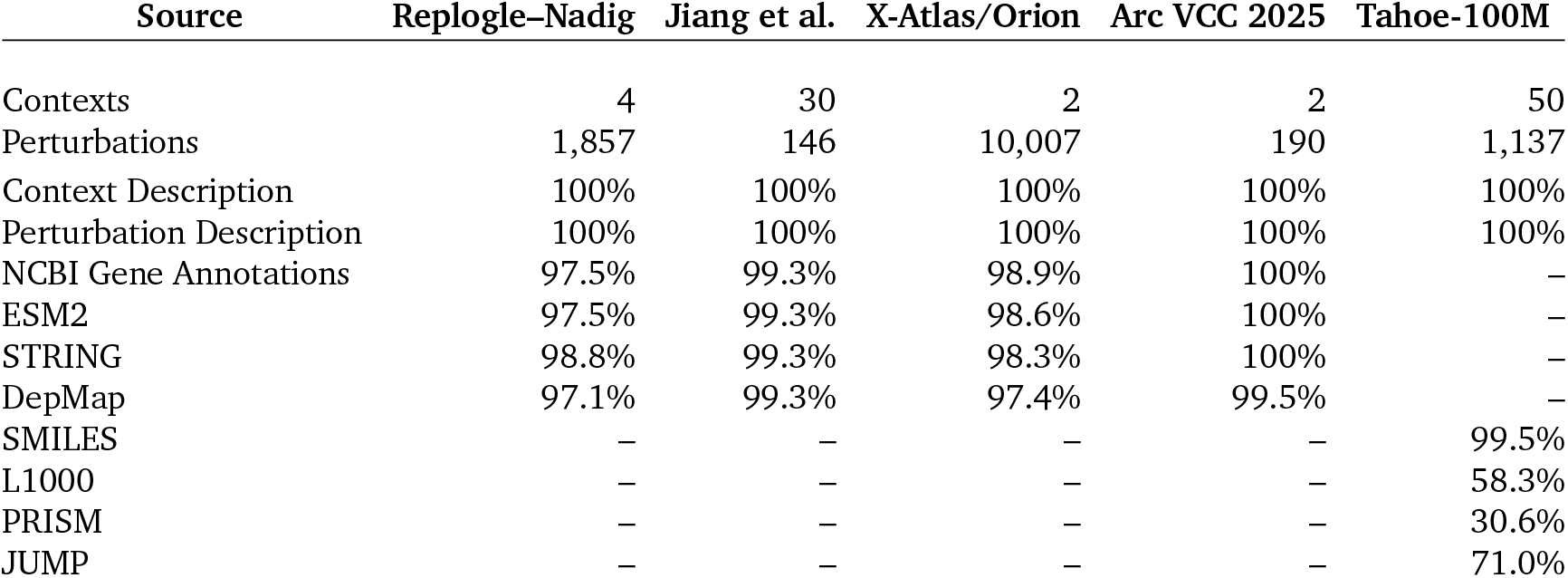
Knowledge Source coverage. Contexts or perturbations for which each source resolves, out of the total in the dataset. Tahoe-100M counts drug–dose perturbations (379 compounds × 3 doses); a compound is covered at all three doses or at none. –, the source does not apply to the dataset’s perturbation type.

### 3.3. Per-gene evidence

The Knowledge Sources describe a context and a perturbation, but not how individual genes behave in them. To give PIE direct access to measured expression, we supplement each gene with per-gene evidence drawn from the data, organized in three channels. Control Mean Evidence is the gene’s mean expression in the control cells of the context, 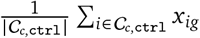, and is available for every context. Same-pert Evidence summarizes the gene’s responses to perturbation *p* in other contexts, that is, over conditions (*c*^′^, *p*) ∈ *T* with *c*^′^ ≠ *c*. Same-context Evidence summarizes its responses to other perturbations in context *c*, over conditions (*c, p*^′^) ∈ *T* with *p*^′^ ≠ *p*. Each response channel records the mean and standard deviation of Δ and ℓ across these conditions, together with the fraction of conditions in which the gene is up- or down-regulated (Appendix A.3). When no training condition contributes to a channel, as for Same-context Evidence in an unseen context, we set the channel to zero. Each statistic is passed through its own small encoder, and a fusion MLP combines the three channels into a single evidence vector **e**_*g*_ per gene.

### 3.4. Encode–process–decode trunk

PIE uses the encode–process–decode design of Perceiver IO (Jaegle et al., 2022), which decouples the size of the input and the output from the size of the model. Learned latent vectors first cross-attend to the Knowledge Source tokens (encode), and the latents then self-attend to one another (process). Finally, gene embeddings query the latents (decode), producing a per-gene feature vector **h**_*g*_ (Appendix A.4). A gene’s query combines what is known about the gene with what has been measured for it: we obtain a single embedding vector of the gene’s NCBI Gene entry, project it to the model width, and add its evidence vector **e**_*g*_. Gene queries do not attend to one another, so each gene is decoded independently. Because the queries are computed from text rather than learned per gene, PIE has no fixed gene vocabulary, and each dataset is decoded on its own set of measured genes.

### 3.5. Training formulation

Three linear heads map **h**_*g*_ to the three labels of Section 3.1, and each head is trained with its own loss. Before each head, **h**_*g*_ is refined with the gene’s evidence vector **e**_*g*_: an MLP takes the two as input and adds its output back to **h**_*g*_ (Appendix Figure 5).

#### DE head

The DE head outputs the probability 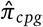 that gene *g* is differentially expressed, which we use to rank genes and threshold at 0.5 to call DE. It is trained with a weighted binary cross-entropy *ℒ*_DE_ over the genes *G*_*cp*_ that pass the CPM filter, where the weights are capped inverse-frequency class weights computed per condition.

#### LFC head

The LFC head outputs a real value 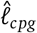, whose sign gives the direction of change. Because the LFC is defined only for the DE genes *D*_*cp*_ ⊆ *G*_*cp*_, we supervise it there, with a Huber loss *H*_*κ*_ (Huber, 1964) on its magnitude and a hinge loss (Cortes and Vapnik, 1995) on its direction:

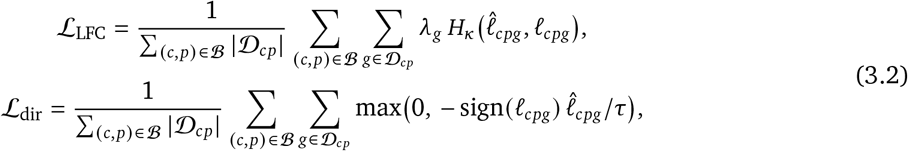

where *λ*_*g*_ down-weights the target gene of a genetic perturbation, *B* is the set of experimental conditions in a minibatch, and *τ* is a fixed scale.

#### Delta-expression head

Rather than regressing Δ_*cpg*_ directly, this head outputs a distribution 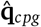 over *K* equal-width bins that tile [−Δ_max_, Δ_max_], with centres *b*_1_ < … < *b*_*K*_, where Δ_max_ is the largest per-dataset 99.9th percentile of |Δ_*cpg*_| over the training set (Appendix A.5). We clip Δ_*cpg*_ to [*b*_1_, *b*_*K*_] and encode it as a two-hot target **t**_*cpg*_ over the neighbouring centres *b*_*k*_ ≤ Δ_*cpg*_ < *b*_*k*+1_, with weight (*b*_*k*+1_ − Δ_*cpg*_)/(*b*_*k*+1_ −*b*_*k*_) on bin *k*, weight (Δ_*cpg*_ −*b*_*k*_)/(*b*_*k*+1_ −*b*_*k*_) on bin *k* + 1, and zero elsewhere. The head is trained with a weighted cross-entropy over all *G* measured genes,

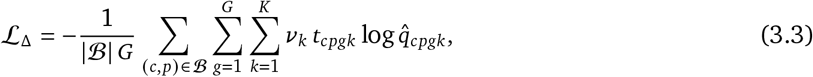

where *v*_*k*_ are capped inverse-frequency bin weights computed per condition. At inference, the predicted delta expression is the centre of the most probable bin, and adding it to the control mean of the context gives the predicted pseudobulk expression of the perturbed cells.

The final loss of the model is a composite loss, the unweighted sum

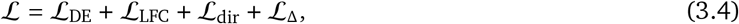

where each term is averaged over the (condition, gene) pairs it supervises in a batch. During training, we randomly set Same-pert and Same-context Evidence to zero for a subset of samples, so that the model learns to predict when either channel is unavailable, as it is for unseen perturbations and contexts.

### 3.6. Generalization to unseen contexts, perturbations, and datasets

PIE represents an unseen context or perturbation in the same way as a seen one, through its Knowledge Sources and the evidence available for it (Appendix A.3). For an unseen context, the training set has no responses in that context, so Same-context Evidence is empty. Likewise, for an unseen perturbation, Same-pert Evidence is empty. For a held-out dataset, Same-context Evidence is empty. Its Same-pert Evidence is present only for perturbations that a training dataset also measured. In every setting, the trained model predicts in a single forward pass, without fine-tuning.

## 4. Experiments

### 4.1. Experimental Setup

#### Datasets

Our main benchmark is the Replogle–Nadig CRISPRi Perturb-seq dataset (Replogle et al., 2022; Nadig et al., 2025), comprising single-gene knockdowns in four cell lines (K562, RPE1, HepG2, and Jurkat). For cross-dataset transfer, we additionally use: (i) Tahoe-100M, a small molecule perturbation dataset (Zhang et al., 2025); (ii) the cytokine-stimulation CRISPRi screen of Jiang et al. (2025); (iii) the X-Atlas/Orion CRISPRi screens in HCT116 and HEK293T (Huang et al., 2025); and (iv) the H1 CRISPRi data from Arc VCC 2025 (the 2025 Virtual Cell Challenge) (Roohani et al., 2025). Appendix A.1 describes dataset preprocessing and reference-label construction.

#### Baselines

We compare PIE to a variety of established methods for context and perturbation generalization. For unseen contexts, we evaluate State, which predicts perturbed cells from control cells (Adduri et al., 2026), Stack, which generates perturbed cells in context from example cells of the same perturbation (Dong et al., 2026), and Rhaister-O, which learns perturbation-specific DE statistics (Svensson et al., 2026). For unseen genetic perturbations in seen contexts, we evaluate COMPASS-N (Liang and Singh, 2026), which transfers responses from nearby perturbations in protein-embedding space, and PRESAGE (Littman et al., 2025), which uses gene knowledge graphs. We also evaluate TabICL (Palla et al., 2026) for both unseen-context and unseen-perturbation prediction, using its task-specific procedures. In our baselines, we further include perturbation mean, which averages a perturbation’s response across training contexts, and context mean, which averages training perturbation responses within a context (Adduri et al., 2026). For an unseen context, context mean predicts the mean expression of its control cells, and is the only applicable baseline when both the context and perturbation are unseen. Appendix B.2 reports baseline implementations.

#### Metrics

We evaluate all models using six metrics drawn from cell-eval (Adduri et al., 2026) and cell-eval2 (Dobin, 2026a): AUPRC, Jaccard score, and accuracy of the predicted direction of change for differentially expressed genes; Spearman correlation between predicted and true log fold changes (LFC) and between predicted and true overall effect sizes; and Discrimination Score using the L1 metric. Together, these evaluate the ability to predict differential expression both in terms of magnitude and direction and ability to discriminate between perturbation responses. Higher values are better for all six metrics. We adapt these metric computations to accept predicted DE probabilities, LFCs, and pseudobulk delta expression directly. All methods are scored on the same test conditions, reference labels, and genes. Appendix B.1 provides the metric definitions, adaptations, and baseline-output conversions.

#### Evaluation Protocol

For held-out contexts, unperturbed control cells are available, but their perturbation responses are excluded from training. PIE’s response-evidence inputs are computed only from training responses, and its parameters are not updated at test time. We select the PIE checkpoint with the highest validation AUPRC; test responses are used only to evaluate predictions. We report mean performance and sample standard deviation across the four folds for within-dataset experiments and across the held-out dataset’s contexts for cross-dataset experiments.

### 4.2. Generalization within dataset

We evaluate three regimes: unseen contexts with seen perturbations, unseen perturbations in seen contexts, and jointly unseen contexts and perturbations. On Replogle–Nadig, each of four folds holds out one test cell line and a subset of perturbations in the remaining three cell lines. Training uses the remaining perturbations in the three non-heldout cell lines. The three test sets isolate context generalization, perturbation generalization, and their combination. One PIE model is trained per fold, and the same selected checkpoint is evaluated in all three regimes. Appendix B.3 gives the full split and training protocols.

PIE achieves the highest score on all metrics for unseen perturbations and on five of six metrics for unseen contexts (Figure 2). For unseen contexts, its AUPRC is 0.424, compared with 0.344 for Rhaister-O, the strongest baseline, and its Spearman LFC is 0.664 versus 0.591. The exception is Spearman effect size, where Rhaister-O scores 0.741 and PIE scores 0.734. For unseen perturbations, PIE has an AUPRC of 0.304 versus 0.094 for TabICL, a greater than threefold improvement. When both entities are unseen, PIE exceeds context mean, the only applicable model, on every metric, reaching an AUPRC of 0.239 versus 0.088 and a Discrimination Score L1 of 0.643 versus 0.502 (effectively random). For context mean, Spearman effect size is undefined because it predicts no change for every perturbation, and Spearman LFC is undefined for an unseen context because it predicts no DE genes. PIE significantly outperforms the strongest baseline on several metrics including AUPRC in similar experiments conducted on Jiang et al. and Tahoe-100M. Appendix B.3 includes these experiment results.

### 4.3. Generalization across datasets

We next test whether PIE can transfer responses to a dataset absent from training. We train on Tahoe-100M, Jiang et al., X-Atlas/Orion, and Arc VCC 2025, and evaluate on the Replogle–Nadig dataset. We exclude from training all cell lines shared with the held-out dataset, including derived cell lines, so this experiment requires generalization to both an unseen dataset and unseen contexts.

We separately evaluate perturbations measured in at least one training dataset and perturbations absent from all training datasets. The latter setting requires simultaneous generalization to an unseen dataset, context, and perturbation. A single checkpoint, selected using validation responses from the training datasets, is evaluated in both regimes. Results are averaged across the four held-out cell lines. Appendix B.4 provides the detailed protocol and additional held-out-dataset evaluations.

For seen perturbations, PIE achieves the highest mean score on five of six metrics (Figure 3). Its AUPRC is 0.378 versus 0.283 for Rhaister-O, the strongest baseline, and its Spearman effect size is 0.627 versus 0.547. The exception is DE direction match, where perturbation mean scores 0.733 and PIE scores 0.727.

For unseen perturbations, PIE exceeds context mean on every defined metric, with an AUPRC of 0.252 versus 0.102 and a Jaccard score of 0.123 versus 0.077. As in the within-dataset experiment, both Spearman correlations are undefined for context mean. Performance declines across all metrics when perturbations are also unseen, but PIE retains predictive signal despite the simultaneous shift in dataset, context, and perturbation. PIE also achieves the highest AUPRC in both regimes when Jiang et al., X-Atlas/Orion or Arc VCC 2025 is held out (Appendix B.4).

### 4.4. Contributions of input signals

We examine the contributions of five input groups: Context Description, Perturbation Knowledge Sources, Control Mean Evidence, Same-context Evidence, and Same-pert Evidence. Using the four Replogle–Nadig folds and three within-dataset regimes, we train separate models for every subset of these groups and compute exact Shapley values: each group’s marginal contribution averaged over subsets of the other inputs. Removed inputs are absent during both training and evaluation. Appendix B.5 gives the full protocol.

The contributions differ substantially across regimes and metrics (Figure 4). For unseen contexts, AUPRC increases from 0.238 without the five input groups to 0.422 with all groups. Same-pert Evidence and Perturbation Knowledge Sources account for 0.169 of this 0.184 gain and also explain most of the improvement in Spearman LFC and Discrimination Score L1. For unseen perturbations, the smaller AUPRC gain of 0.045 is distributed across all five groups. In contrast, Perturbation Knowledge Sources account for nearly the entire gain in Discrimination Score L1, contributing 0.205 of 0.207. These inputs therefore contribute more strongly to distinguishing perturbation responses than their AUPRC contribution alone suggests.

When both entities are unseen, the total AUPRC gain is only 0.010. Perturbation Knowledge Sources have a negative mean AUPRC attribution, but account for most of the improvement in DE direction match, Spearman LFC, and Discrimination Score L1. Thus, their contribution depends on the prediction target: improvements in response direction, magnitude, and specificity do not necessarily translate into improved DE ranking. Appendix B.5 further separates the contributions of the individual Perturbation Knowledge Sources.

## 5. Conclusion

We introduced PIE, a perturbation effect prediction model, and demonstrated its ability to generalize to unseen cellular contexts, perturbations, and experimental datasets. PIE incorporates external knowledge sources, control expression, and measured responses from observed conditions, providing a richer representation of each experimental condition than transcriptomic data alone. To provide a more direct learning signal for perturbation effects, PIE simplifies the conventional task of predicting cell-level expression states by instead focusing on population-level responses.

PIE outperformed baselines specialized for unseen-context, unseen-perturbation, or unseen-dataset generalization, and retained predictive signal when multiple axes were jointly unseen. Ablations revealed that the contributions of its inputs differ substantially across generalization settings and evaluation metrics, highlighting the importance of representing different sources of information for different generalization tasks. While PIE improved generalization over existing methods, performance declined when contexts and perturbations were jointly unseen. Performance was also weaker for perturbations with small effect sizes, where distinguishing biological signal from experimental variation is particularly challenging.

Our results suggest that further model improvements will require perturbation data spanning more diverse experimental conditions, as well as better measurements of perturbation effects within those conditions. We intend PIE to serve as an extensible framework for integrating these advances and guiding future model development and data collection. As perturbation datasets grow in size and diversity and experimental technologies improve, more granular measurements that capture heterogeneity within cell populations may provide additional training signal for predicting perturbation effects. Richer characterization of the biological system being perturbed, including its spatial and temporal context, may further improve generalization across experimental conditions.

## Code and Data Availability

All code for this project is available at:

https://github.com/ArcInstitute/pie

Model checkpoints and Data are available at:

https://huggingface.co/collections/arcinstitute/pie

## Acknowledgements

We thank Chiara Ricci-Tam, Brian Plosky and Joseph Caputo for their support in drafting this manuscript. We thank Alexander Dobin, Christopher Carpenter, Jeremy Sullivan, Reshma Mehta, Thomas Sandman, Dara Leto, Tony Hua and Sadie VanHorn for their support and feedback on this work. Furthermore, we thank Usman Muzaffar, Silvana Konermann, Megan van Overbeek and the leadership at Arc Institute for their guidance on this work. This study was supported by Arc Institute. We also thank the community for producing large scale perturbation and annotation data used for training such models.

## AI use statement

In this work, we used generative AI tools to help develop conceptual frameworks, propose and refine hypotheses, design and provide feedback on research methodology and experiments, implement methods, clean and reformat datasets, support qualitative and thematic data analysis, and interpret results. We have not used generative AI tools to generate synthetic datasets. Formulating mathematical claims, providing critical ingredients for proving mathematical claims, assisting in the writing of proofs, and assisting with translation are not applicable to this work. Additionally, we used generative AI tools to create and modify scientific figures, suggest experimental parameters, create and edit software code, draft parts of the paper, summarize or analyse existing literature, discover research topics or identify gaps, brainstorm, source and search for information, edit the paper to improve readability, identify relevant literature, format references, suggest a structure for the paper, and propose its title and keywords.

AI coding agents wrote all project code, including the model, the training and evaluation pipelines, and data preprocessing, under the direction and supervision of the authors. AI agents also executed experiments, orchestrating some of them autonomously under the same direction and supervision. AI tools helped design the templates that render database records as text for the Context Description, Perturbation Description, and NCBI Gene Annotations Knowledge Sources (Appendix A.2). The authors reviewed the code alongside AI review agents, and reviewed all results, hypotheses, and literature produced with AI assistance. We have reviewed all AI-assisted work and take responsibility for the final content of this work, including text, claims or artifacts produced with the aid of generative AI.

## Appendix

### A. Method details

#### A.1. Datasets and Preprocessing

Every dataset starts from raw UMI counts. For genetic perturbation datasets, we apply the on-target knockdown filter of cell-load (Adduri et al., 2026) in each context. It keeps perturbations that reduce their target gene below 30% of its control expression, keeps the cells within them that reduce it below 50%, and drops perturbations left with fewer than 30 cells. Control cells are never filtered. In every dataset, genetic or chemical, counts are then scaled to 10^4^ per cell (CP10K), giving the normalized expression *n*_*ig*_ of gene *g* in cell *i*, and log-transformed, *x*_*ig*_ = log (1 +*n*_*ig*_), which gives the log-normalized expression.

- *Replogle–Nadig* combines the CRISPRi essential-gene screens of Replogle et al. (2022) in K562 and hTERT-RPE1 with those of Nadig et al. (2025) in HepG2 and Jurkat. We use raw counts restricted to the 6,642 genes shared by the four public releases (figshare for Replogle et al., 2022; GEO GSE264667 for Nadig et al., 2025), in which the knockdown filter had already been applied. The data comprise 5,459 experimental conditions over 4 contexts.
- *Tahoe-100M* (Zhang et al., 2025) profiles 379 compounds, each at 0.05, 0.5, and 5 μM, in 50 cancer cell lines, with DMSO-treated control cells per line. We keep the 95.6M of 100.6M cells that pass the release’s cell filter, on its 18,151 genes, giving 56,827 experimental conditions.
- *Jiang et al*. (2025) is a CRISPRi screen of 218 signalling regulators in 6 cell lines, each under one of 5 cytokine stimuli, giving 30 contexts. We take raw counts from the five pathway objects on GEO (GSE281048) and concatenate them on their 33,056 shared genes. The knockdown filter keeps 461 of 1,626 experimental conditions.
- *X-Atlas/Orion* (Huang et al., 2025) is a genome-wide dual-guide CRISPRi screen in HCT116 and HEK293T, downloaded from Figshare+(doi:10.25452/figshare.plus.29190726). We keep the 37,443 genes detected in both lines and, following the release paper, the perturbations with at least 25 cells, at least 30% mean knockdown, and an expressed target. The knockdown filter then keeps 13,284 experimental conditions.
- *Arc VCC 2025* is the H1 hESC CRISPRi data of the 2025 Virtual Cell Challenge (Roohani et al., 2025), on 18,151 genes. Its training and validation files are two contexts, each with its own copy of the control pool. The knockdown filter keeps 146 of 150 and 44 of 50 perturbations, giving 190 experimental conditions.

##### DE calls

DE is called with gpudge (Dobin, 2026b), separately for each context against its own control cells. For each (perturbation, gene), a two-sided Wilcoxon rank-sum (Mann–Whitney *U*) test compares the perturbed and control cells. A gene is tested only when its mean CPM exceeds 5 in the perturbed or in the control group; we say it *passes the CPM filter*. The *p*-values are Benjamini–Hochberg corrected per perturbation across the tested genes, and a gene is DE, *y*_*cpg*_ = 1, when the adjusted value is below 0.05. No fold-change criterion enters the call. An untested gene is not DE.

##### LFC

The LFC target is the log2 fold change that gpudge reports with each test. With the arithmetic mean CP10K 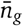 expression over a set of cells,

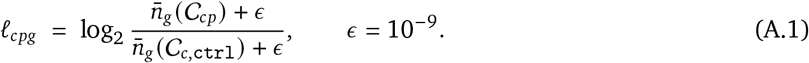

It is defined for DE genes with a finite fold change and is not clipped. A DE gene is up-regulated when ℓ_*cpg*_ > 0 and down-regulated otherwise.

##### Pseudobulk delta expression

For context *c*, perturbation *p*, and gene *g*,

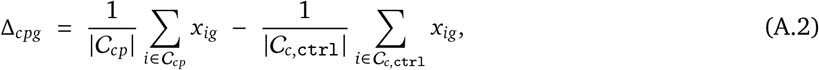

where *x*_*ig*_ is the log-normalized expression of cell *i, C*_*cp*_ the perturbed cells, and *C*_*c*, ctrl_ the control cells of context *c*. It is defined for every measured gene, whether or not the gene passes the CPM filter.

##### Measured genes

When a run trains on several datasets, each experimental condition is decoded and supervised only on the *G* genes of its own dataset.

#### A.2. Knowledge Sources

##### A.2.1. Sources

###### Context Knowledge Sources

The *Context Description* is the only context source. For a cell line, it is rendered from the Cellosaurus entry of the line (Bairoch, 2018):

~~~
Context Name: {context} ;
Recommended Name: {name} ;
Synonyms: {synonyms} ;
Cell Line Category: {category} ;
Organism: {organism} ;
Sex: {sex} ;
Age: {age} ;
Population: {population} ;
Disease: {disease} ;
Derived From Site: {site} ;
Cell Type: {cell type} ;
Parent Cell Line: {parent} ;
Genetic Integration: {label} via {method} ;
Sequence Variation: {genes}: {type}, {description} ({zygosity}; {note})
~~~

For Jiang et al., where a context is a cell line under a stimulus, the Cellosaurus entry is followed by a record of the stimulus. We derived one record for each of the five stimuli from the UniProt entry of its ligand (The UniProt Consortium et al., 2024), giving its name, family, receptors, canonical signalling pathway, and a short description:

~~~
{Cellosaurus entry} ;
Stimulation: {name} ({abbreviation}) ;
Stimulation Family: {family} ;
Stimulation Receptors: {receptors} ;
Stimulation Signaling: {pathway} ;
Stimulation Description: {description}
~~~

The rendered text is embedded with `text-embedding-3-large` (OpenAI, 2024) into one 3,072-dimensional vector. A text longer than the model’s 8,191-token limit is split into windows with 50% overlap, and the window embeddings are averaged.

##### Genetic Perturbation Knowledge Sources

- The *Perturbation Description* is rendered from the NCBI Gene entry of the target gene (Brown et al., 2015):

~~~
Gene Name: {perturbation} ;
HGNC Gene Symbol: {symbol} ;
Full Name: {full name} ;
Synonyms: {synonyms} ;
Gene Type: {type} ;
Organism: {organism} ;
Lineage: {lineage} ;
Map Location: {cytoband} ;
Alt Designations: {designations} ;
Gene Summary: {summary}
~~~ It is embedded like the Context Description.
- *NCBI Gene Annotations* list, in full, the Gene Ontology terms (Ashburner et al., 2000; The Gene Ontology Consortium et al., 2025), OMIM disease associations (McKusick-Nathans Institute of Genetic Medicine, Johns Hopkins University, 2026), and gene-group memberships of the NCBI Gene record. GO terms are sorted by evidence count and diseases by confidence. The gene summary is left out, since the Perturbation Description carries it:

~~~
Gene: {symbol}
=== GO BIOLOGICAL PROCESS ===
Gene Ontology biological process annotations, sorted by evidence count.
- {GO ID}: {term} (evidence: {count})
=== GO MOLECULAR FUNCTION ===
(same layout)
=== GO CELLULAR COMPONENT ===
(same layout)
=== DISEASE ASSOCIATIONS ===
Known disease associations with confidence scores and OMIM (Online Mendelian Inheritance in Man) identifiers.
-{disease} (confidence: {score}, MIM: {MIM ID})
=== GENE GROUPS ===
Gene family and group memberships by relationship type.
-{relationship}: {members}
~~~ An empty section reads “None”. The text is encoded by `Qwen3.5-35B-A3B-Base` (Qwen Team, 2026), and the final hidden state of every token is kept, one 2,048-dimensional vector per token. This is the only per-token source. Texts run from 122 to 5,874 tokens, below the 8,192-token limit.
- *ESM2* (Lin et al., 2023): the `esm2_t33_650M_UR50D` embedding of the protein sequence, mean-pooled over residues. Sequences longer than 1,022 residues are embedded in windows of 1,022 with stride 511.
- *STRING* (Szklarczyk et al., 2025): the 512-dimensional SPACE embedding (Hu et al., 2025) of the STRING v12.0 protein network.
- *DepMap* (DepMap, Broad, 2025; Dempster et al., 2021): the Chronos gene effect (release 25Q3) across 1,186 cell lines, with missing values set to 0.

##### Chemical Perturbation Knowledge Sources

- The *Perturbation Description* is rendered from the PubChem entry of the compound (Kim et al., 2025), followed by the dose:

~~~
Compound Name: {name} ;
Synonyms: {first 10 synonyms} ;
Molecular Formula: {formula} ;
Molecular Weight: {weight} g/mol ;
SMILES: {SMILES} ;
Connectivity SMILES: {connectivity SMILES} ;
XLogP: {XLogP} ;
Topological Polar Surface Area: {TPSA} square angstroms ;
Complexity: {complexity} ;
Formal Charge: {charge} ;
Hydrogen Bond Donors: {donors} ;
Hydrogen Bond Acceptors: {acceptors} ;
Rotatable Bonds: {bonds} ;
Compound Description: {description} ;
Dose: {dose} micromolar
~~~ It is embedded like the Context Description.
- *SMILES* (Ahmad et al., 2022): the `ChemBERTa-77M-MTR` embedding of the SMILES string, mean-pooled over tokens (384 dimensions).
- *L1000* (Subramanian et al., 2017): the transcriptional activity score of the compound across 83 cell lines (GEO GSE92742 and GSE70138). For each line, the signature nearest the perturbation’s dose and a 24 h time point is used.
- *PRISM* (Corsello et al., 2020): secondary-screen log_2_ viability across 480 cell lines, at the measured dose nearest the perturbation’s.
- *JUMP* (Chandrasekaran et al., 2023): 737 processed Cell Painting features. JUMP does not vary the dose, so every dose of a compound receives the same vector.

The three profiles enter unstandardized, and a cell line without a measurement is set to 0. No source contains measurements from the datasets of Appendix A.1. DepMap includes some of the same cell lines, such as K562, but from independent CRISPR screens.

##### A.2.2. Integration into PIE

Let source *i* be present for an experimental condition with *m*_*i*_ vectors of width *d*_*i*_. Each vector passes through an RMSNorm over *d*_*i*_ and a bias-free linear map to the model width *d* = 768. A learned source-type embedding, indexed from a fixed registry of source names, is added to every token of the source, with element-wise dropout of 0.05, and a per-source RMSNorm follows. The present sources are concatenated into one key-value set of *M* = ∑_*i*_*m*_*i*_ tokens, with no positional encoding, so an absent source contributes no tokens. Within a batch, each source is zero-padded to its longest sequence, and cross-attention masks the padded positions. A condition for which no source resolves receives a single all-zero carrier token, so the key-value set is never empty. A run uses the sources that fit its perturbations: the four gene sources for Replogle–Nadig and Jiang et al., the four compound sources for Tahoe-100M, and all ten in the cross-dataset recipe, each with both descriptions.

#### A.3. Per-gene evidence

##### Channels

Each response statistic below comes with a presence flag and the log number of contributing responses. Delta expression and LFC are divided by a global training standard deviation.

- *Control Mean Evidence*: the gene’s mean log-normalized expression over control cells of the context, given as the value, its log, and a not-expressed flag. It is always present.
- *Same-pert Evidence*: over the training responses to the same perturbation, the mean and standard deviation of the gene’s pseudobulk delta expression and of its LFC, and the fractions of tested responses in which it is up- or down-regulated. In the cross-dataset recipe, a provenance block also records which training datasets measured the perturbation. The channel is empty for an unseen perturbation, and for a held-out dataset unless another dataset measured the perturbation.
- *Same-context Evidence*: the same statistics over the training responses in the same context. It is empty for an unseen context and for a held-out dataset.

##### Donors

All response evidence comes from the training set.

##### Fusion

Each statistic has its own small encoder. The output of a response encoder is multiplied by its presence flag, so an empty channel contributes exactly zero. The encoded blocks are concatenated and fused by an MLP into a 128-dimensional evidence vector per gene (Figure 5). The vector is added to the gene’s query and conditions the task adapter before each head (Figure 1).

**Figure 1.**
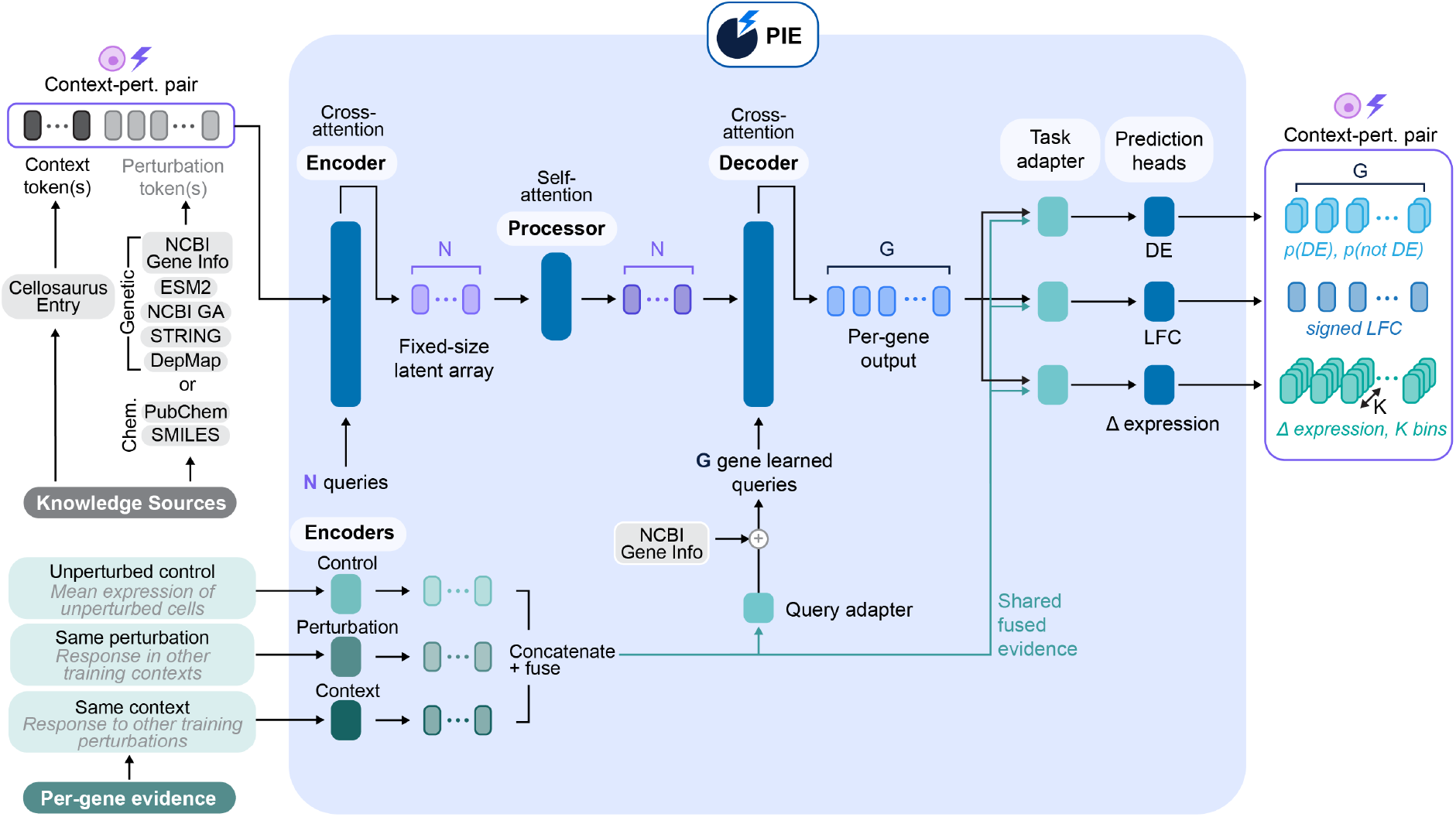
PIE overview. A sample is a (context, perturbation) pair. Knowledge Sources describe contexts and perturbations with variable-size sets of embeddings from external databases. A source with no entry for a context or perturbation contributes no tokens. Learned latents query these tokens (Encoder) and subsequently are processed by a bidirectional transformer (Processor). Each measured gene is represented by a text embedding derived from its NCBI Gene record, and is further augmented with experimental evidence, such as the control-cell mean expression of the context, summary statistics of responses to the same perturbation in other contexts, and summary statistics of responses to other perturbations in the same context. These response statistics are derived exclusively from the training set. In the Decoder, each gene queries the latents, producing a per-gene feature vector. Three heads, each preceded by a task adapter, predict a binary DE call, a signed log fold change, and a binned pseudobulk delta expression, for every gene.

**Figure 2.**
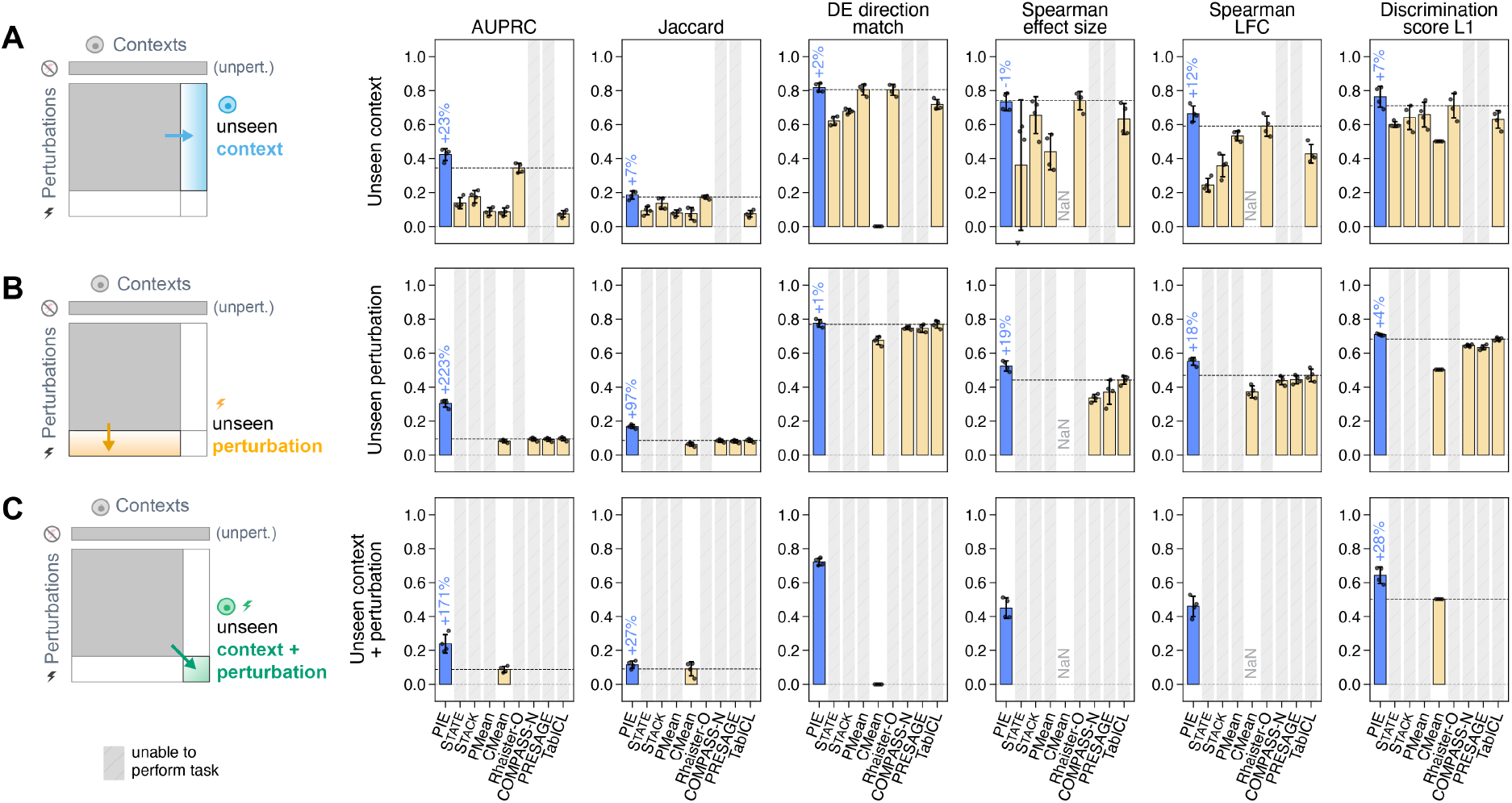
Generalization within the Replogle–Nadig dataset. The three generalization regimes. (A) Unseen contexts with seen perturbations. (B) Unseen perturbations in seen contexts. (C) Both unseen. Each panel pairs a regime schematic (left) with its scores (right). Bars show means, and error bars show standard deviations across four folds, each holding out one cell line. In the score panels, gray-shaded slots mark models that cannot perform the task, and the dashed line marks the strongest baseline. Higher is better for all metrics. PIE has the highest AUPRC in all three regimes, exceeding the strongest baseline by 0.08 for unseen contexts, 0.21 for unseen perturbations, and 0.15 for both unseen.

**Figure 3.**
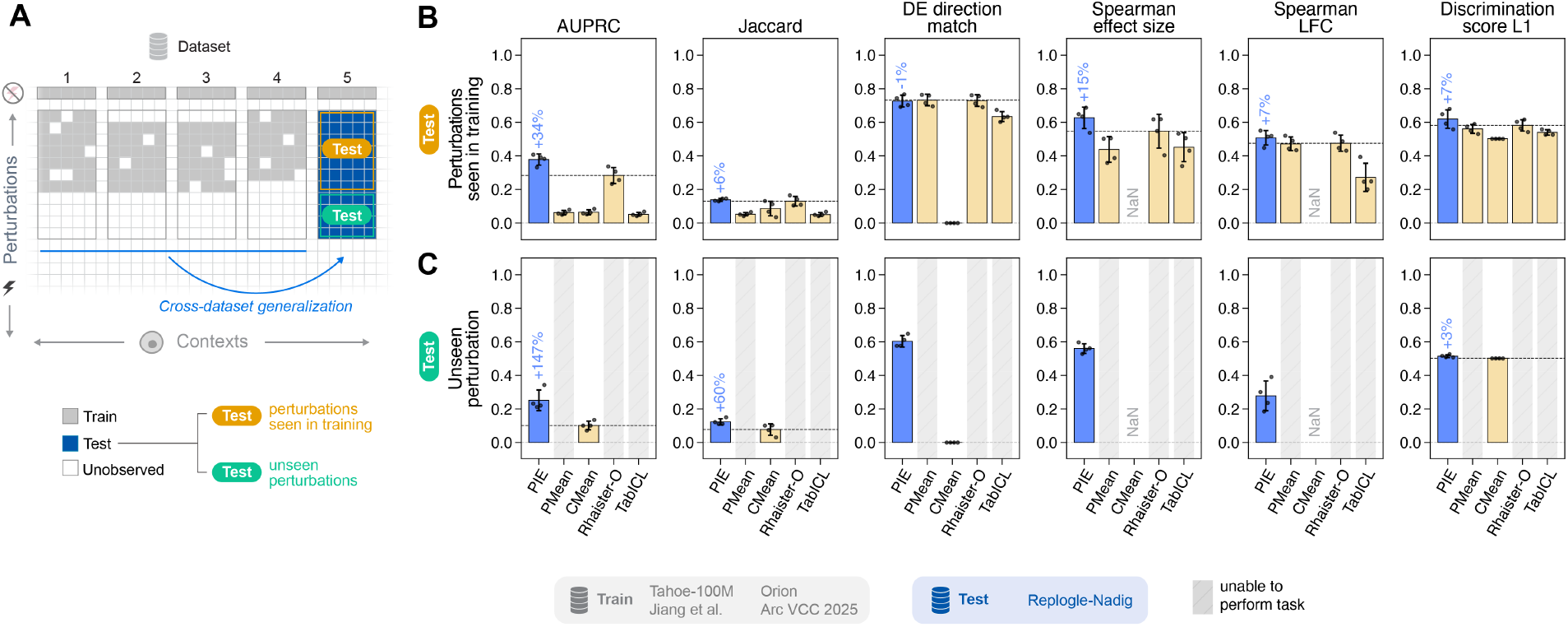
Generalization to an unseen dataset, Replogle–Nadig. (A) Training and test schematic: models trained on four other datasets predict responses in the four Replogle–Nadig cell lines. (B) Scores for perturbations seen in training. (C) Scores for perturbations unseen in training, requiring zero-shot generalization to an unseen dataset, context, and perturbation. Bars show means, and error bars show standard deviations across the four cell lines. In the score panels, gray-shaded slots mark models that cannot perform the task, and the dashed line marks the strongest baseline. Higher is better for all metrics. PIE has the highest AUPRC in both regimes, exceeding the strongest baseline by 0.10 for seen perturbations and 0.15 for unseen perturbations.

**Figure 4.**
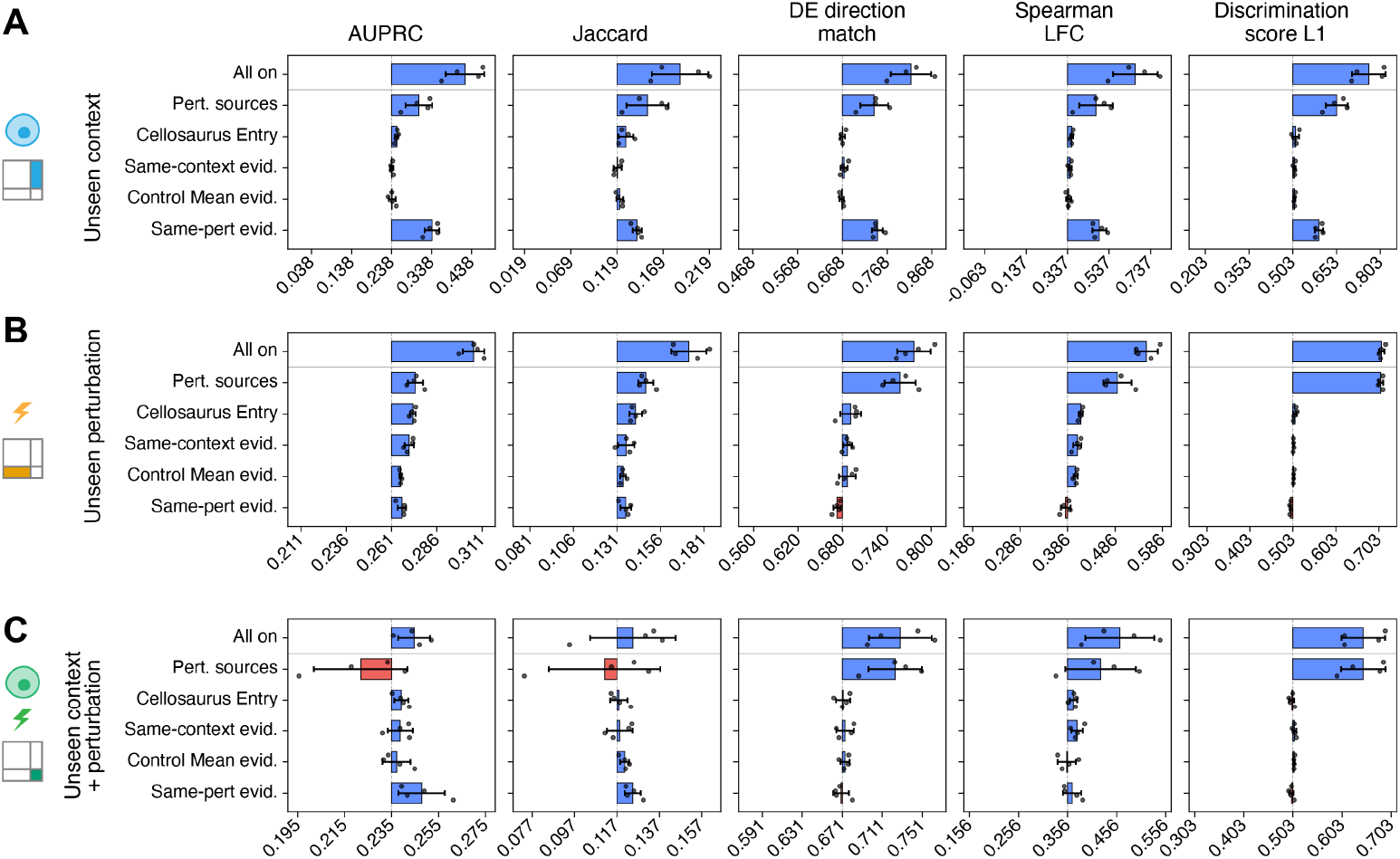
Contribution of input signals. Shapley values of five input groups on the within-dataset task, computed from models trained with every subset of the groups: Context Description (Cellosaurus entry), Perturbation Knowledge Sources (Pert. sources), Control Mean Evidence, Same-context Evidence, and Same-pert Evidence. (A), (B), and (C) are the three generalization regimes. Each axis is centered (dashed line) on the model without any group, and the top bar (All on) is the difference between the models with all and with no groups; tick labels are metric values. Bars show means and std across four folds, each holding out one cell line; dots are folds. Same-pert Evidence and the Perturbation Knowledge Sources account for most of the AUPRC gain for unseen contexts.

**Figure 5.**
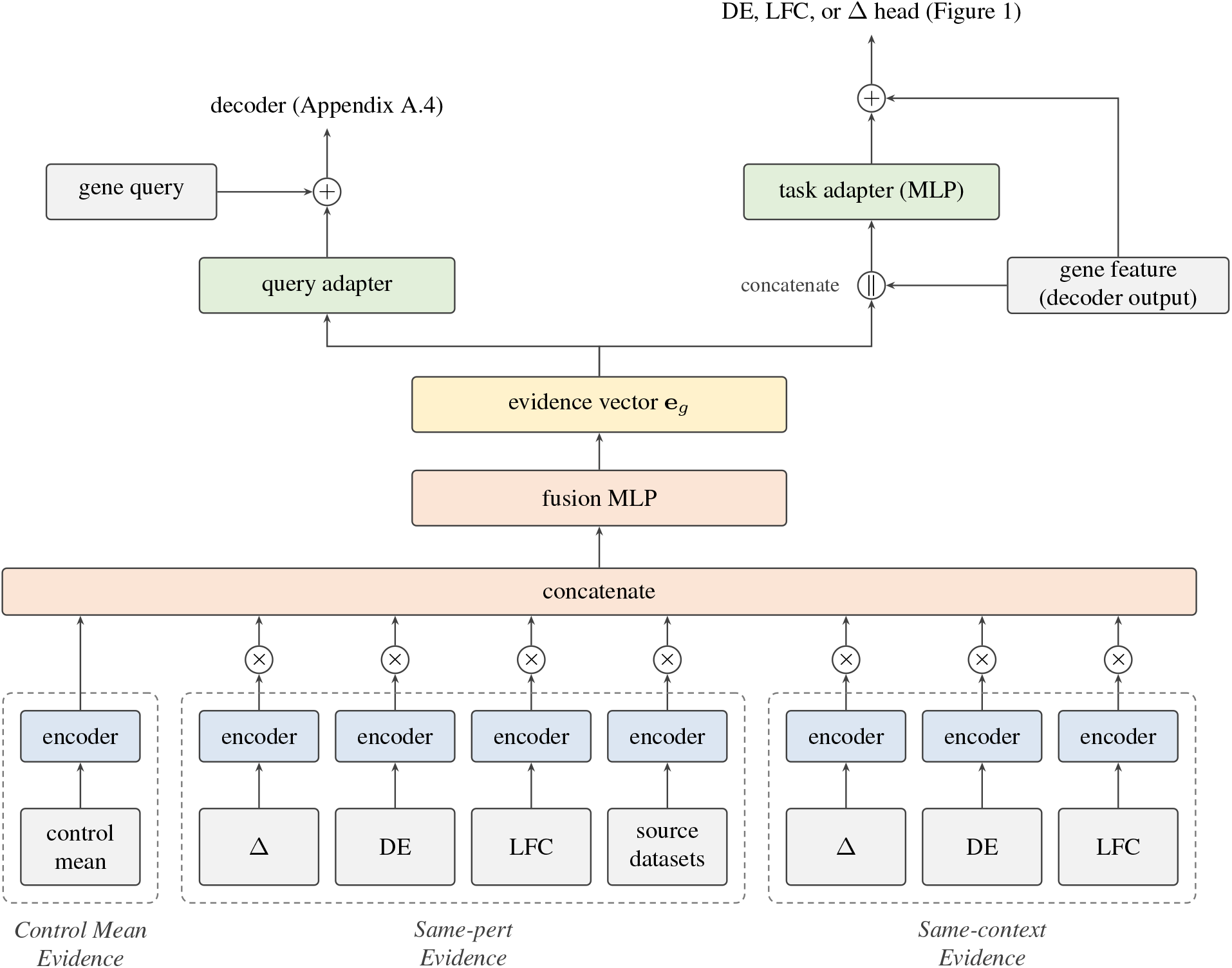
Evidence fusion. Each block of per-gene evidence is encoded separately, and the response blocks are gated by their presence flags. The concatenated blocks are fused into the evidence vector **e**_*g*_, which a query adapter adds to the gene query and task adapters pass to the three heads.

#### A.4. Encode–process–decode trunk

##### Blocks

The trunk has three stages (Figure 6). A learned array of latents cross-attends the Knowledge Source tokens, the latents then pass through self-attention blocks, and the gene queries cross-attend the latents. Each stage stacks several blocks. Every block applies attention and a two-layer GELU feed-forward (Hendrycks and Gimpel, 2016), each pre-normalized with RMSNorm (Zhang and Sennrich, 2019) and wrapped in a residual connection. Attention applies query–key normalization with a learned per-head temperature (Henry et al., 2020). Cross-attention also normalizes its keys and values. Gene queries do not attend to each other, so each gene is decoded independently of the others.

**Figure 6.**
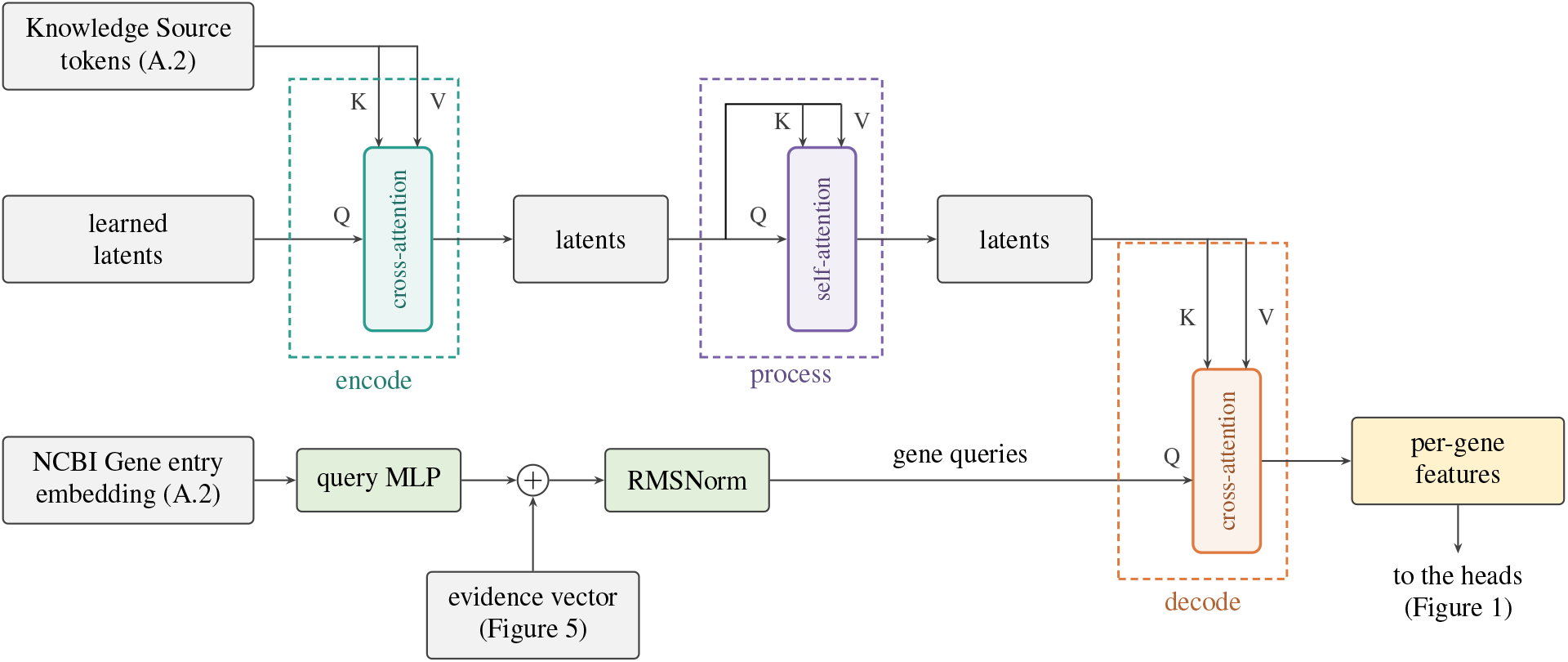
Encode–process–decode trunk. Following Perceiver IO (Jaegle et al., 2022), learned latents cross-attend the Knowledge Source tokens (encode), attend to one another (process), and are read out through cross-attention by one query per gene (decode). Each gene query is the gene’s embedded NCBI Gene entry, passed through a query MLP, summed with its evidence vector, and normalized.

##### Gene queries

A gene’s query is built from its NCBI Gene entry, the same text as the Perturbation Description of a genetic perturbation (Section 3.2; Appendix A.2), embedded with text-embedding-3-large. A query MLP maps the embedding to the model width, the query adapter adds the gene’s evidence vector (Appendix A.3), and an RMSNorm follows. The Perturbation Description has its own projection, separate from the query MLP. Because a query depends only on the gene’s text, there is no learned per-gene parameter, and the gene vocabulary can differ across datasets. Both in training and at inference, each experimental condition is decoded on the full gene vocabulary of its dataset.

#### A.5 Training formulation

Each head is a linear layer applied to **h**_*g*_ +*a* (**e**_*g*_, **h**_*g*_), where **h**_*g*_ is the decoded gene feature, **e**_*g*_ the evidence vector, and *a* a head-specific MLP on their concatenation (Figure 5). As in Section 3.5, *G*_*cp*_ is the set of genes of an experimental condition (*c, p*) that pass the CPM filter (Appendix A.1), *D*_*cp*_ ⊆ *G*_*cp*_ its DE genes, and *G* the number of measured genes. We drop the subscript *c p* from per-gene quantities.

##### DE head

The label is the binary DE call *y*_*g*_ of Appendix A.1. The head outputs two logits, and the DE probability 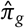 is the softmax probability of the DE class; a gene is called DE when 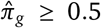. L_DE_ is the cross-entropy over G_*cp*_, with each gene weighted by the inverse frequency of its class in the condition.

##### LFC head

The label is the log_2_ fold change ℓ_*g*_ of Appendix A.1, defined for DE genes. The head outputs one value 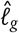, whose sign gives the predicted direction. *ℒ*_LFC_ and *ℒ*_dir_ are given in Equation 3.2. *ℒ*_LFC_ is the Huber loss *H*_*κ*_ with *κ* = 1 between 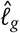 and ℓ_*g*_ over *D*_*cp*_; for a genetic perturbation, the target gene’s term is down-weighted. The direction loss is a hinge loss (Cortes and Vapnik, 1995) with zero margin on the sign of the prediction; it is zero when 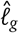 has the sign of ℓ_*g*_.

##### Delta-expression head

The label is the pseudobulk delta expression Δ_*g*_ of Appendix A.1, discretized on a grid fixed once per run from its training rows. For each training dataset *s*, let *R*_*s*_ be the 99.9th percentile of |Δ| and *φ*_*s*_ its minimum fold change, a resolution constant of 1.1 for Replogle–Nadig and Tahoe-100M and 1.01 for the other datasets. With Δ_max_ = max_*s*_ *R*_*s*_ and *w* = min_*s*_ ln *φ*_*s*_, the grid has the smallest odd number of bins *K*≥ 2Δ_max_ / *w* over [−Δ_max_, Δ_max_], so one bin is centred on zero; values beyond ±Δ_max_ are clamped. The target is two-hot: its mass is split between the two bins whose centres bracket the value, so that the expected centre equals the value. The head outputs *K* logits, and the predicted delta expression is the centre of the most probable bin. Adding the predicted delta expression to the control mean gives the predicted pseudobulk expression. *ℒ*_Δ_ is the cross-entropy against the two-hot target over the *G* measured genes with a finite Δ_*g*_, with each gene weighted by the inverse frequency of its target bin in the condition. The grid is computed from training data only, and validation and test values are binned on this same grid.

##### Training

Parameters are trained from random initialization with AdamW (Loshchilov and Hutter, 2019), with a learning rate that warms up linearly, stays constant, and then decays on a cosine schedule. The Knowledge Source embeddings are precomputed and fixed. Each response-evidence channel is emptied, independently, for a random subset of experimental conditions. The checkpoint with the best validation AUPRC is kept.

#### A.6. Hyperparameters

The model and training hyperparameters for the within-dataset and cross-dataset experiments are summarized in Table 2.

**Table 2.** Hyperparameters. Shared by the within-dataset (Section 4.2) and cross-dataset (Section 4.3) runs except where noted.

| Group | Setting | Value |
| --- | --- | --- |
| Trunk | model width $d$ | 768 |
|  | latents | 512 |
|  | attention heads (width) | 8 (96) |
|  | encoder / process / decoder blocks | 4 / 6 / 4 |
|  | feed-forward width | 1,536 |
|  | dropout | 0.1 |
| Knowledge Sources | source-type embedding dropout | 0.05 |
| Evidence | per-block encoder width | 64 |
|  | evidence vector width | 128 |
|  | fusion dropout | 0.1 |
|  | response-channel dropout | 0.25 |
| Losses | DE class-weight cap | 10 |
|  | LFC Huber threshold | 1 |
|  | target-gene LFC weight | 0.001 |
| | direction scale $\tau$ | 0.25 |
|  | delta-expression bin-weight cap | 5 |
|  | delta-expression range percentile | 99.9 |
| Optimization | optimizer | AdamW, $\beta = (0.9, 0.999)$ |
|  | training steps | 5e3 within dataset; 1e4 cross dataset |
| | learning rate | $10^{-4}$ |
|  | weight decay | 0.01 |
| | schedule | 5% linear warmup, 4,250 steps at peak, cosine decay to $10^{-6}$ over the rest |
|  | effective batch size | 128 |
|  | precision | bf16 mixed |
|  | gradient clipping | 10 |

### B. Experiment details

#### B.1. Evaluation metrics

##### Notation

For a test experimental condition (*c, p*) of context *c* and perturbation *p*, dropping the subscript *c p*, let *D* be the reference DE genes among the *G* genes measured in the test dataset, and ℓ_*g*_ and Δ_*g*_ the reference LFC and pseudobulk delta expression of gene *g* (Appendix A.1). A model predicts a DE probability 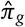, an LFC 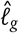, and a pseudobulk delta expression 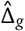 for every measured gene, and its predicted DE set is 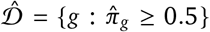. For a genetic perturbation, *D*^−^ and 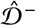 denote these sets without the target gene of *p*; otherwise they equal *D* and 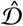.

##### Metrics

We report performance on the following metrics:

- **AUPRC** is the average precision of the measured genes ranked by 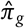, with the genes in *D* as positives. An experimental condition whose genes are all positive or all negative scores 0.
- **Jaccard score** is 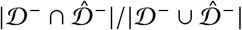, and 1 when the union is empty.
- **DE direction match** is the fraction of genes in D whose 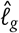 has the sign of ℓ_*g*_, and 0 when *D* is empty. It does not depend on 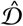.
- **Spearman effect size** is the Spearman correlation between |*D*| and 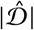 across the perturbations of a context with |*D*| > 0.
- **Spearman LFC** is the Spearman correlation between 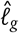 and ℓ_*g*_ over the genes in *D*.
- **Discrimination Score L1**. Let *N* be the number of perturbations in the context and *r*_*p*_ the number of other perturbations *q* whose reference profile Δ^(*q*)^ is closer in L1 distance to the prediction 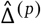 than *p*’s own reference Δ^(*p*)^ is. For a genetic perturbation, the target gene of *p* is removed from all profiles. The score is 1 − *r*_*p*_ /*N*, so 1 is perfect and random profiles score about 0.5. Predicted deltas are clipped so that control expression plus delta is nonnegative.

##### Aggregation

Spearman effect size is computed once per context; the other five metrics are computed per experimental condition and averaged over the perturbations of a context. Each context thus contributes one value per metric, and we report the unweighted mean over contexts. A Spearman correlation is undefined when fewer than two values enter it or either side is constant; such values are excluded from the means.

##### Adaptation

cell-eval and cell-eval2 score single-cell predictions. They run a Wilcoxon rank-sum test of predicted perturbed cells against control cells, call a gene DE at Benjamini–Hochberg adjusted *p* < 0.05, rank genes by −log_10_ adjusted *p* for AUPRC, take LFCs from the test, and compute the Discrimination Score on pseudobulk expression. Our adaptation replaces the predicted significance with 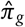: DE calls are 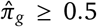, and AUPRC ranks by 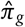. The LFC metrics take 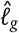, and the Discrimination Score takes 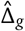. Otherwise the metrics follow the v2 definitions of cell-eval2, including its values for degenerate experimental conditions, with one exception: an undefined Spearman correlation is excluded from the mean instead of scoring −1. Discrimination Score L1 uses the cell-eval implementation of Adduri et al. (2026).

##### Rescoring baselines

Each baseline’s output is converted to 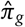, 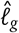 and 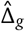 and scored by the same code as PIE.

- *Single cells* (State; Stack; TabICL for unseen contexts). We run gpudge (Dobin, 2026b), with the settings used for the reference labels, on the predicted perturbed cells against the control cells of the model’s output, which for State are its own predicted controls and for Stack its synthetic controls. This gives an adjusted *p*-value *q*_*g*_ and 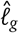. We set 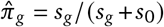 with *s*_*g*_ = −log_10_ max (*q*_*g*_, 10^−10^) and *s*^0^ = −log_10_ 0.05. 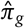 decreases with *q*_*g*_: *q*_*g*_ = 0.05 gives 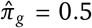, *q*_*g*_ < 0.05 gives 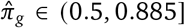, and *q*_*g*_ > 0.05 gives 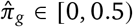, with 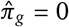 at *q*_*g*_ = 1. The floor of 10^−10^ on *q*_*g*_ sets the upper bound of 0.885; without it, *q*_*g*_ = 0 would make *s*_*g*_ infinite and 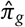 undefined. A gene is therefore called DE 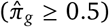 exactly when *q*_*g*_ ≤ 0.05, the threshold of the reference labels. 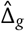 is the difference in mean log-normalized expression between predicted perturbed and control cells. Genes removed by the gpudge expression filter receive *q*_*g*_ = 1 and 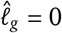.
- *Pseudobulk shifts* (perturbation mean, context mean, COMPASS-N, PRESAGE; TabICL for unseen perturbations). We draw 500 control cells of the test context once without replacement and add the predicted shift to each, clipping at zero. The resulting cells and the 500 controls are then converted as single cells.
- *DE statistics* (Rhaister-O). Rhaister-O predicts an adjusted *p*-value, an LFC, and a delta for each gene. The adjusted *p*-value is mapped to 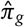 as above, and the other two are used directly.

For an unseen context, context mean predicts a zero shift. It therefore calls no gene DE, predicts 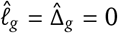, and leaves both Spearman correlations undefined.

#### B.2. Baselines

Each baseline is run with its authors’ code where available, and its predictions are converted and scored as in Appendix B.1.

- *State* (Adduri et al., 2026) is a transformer over sets of cells, trained with an energy-distance loss, that maps the control cells of a context to perturbed cells. It represents each perturbation by a learned embedding and so applies only to perturbations seen in training.
- *Stack* (Dong et al., 2026) is a single-cell foundation model that predicts a perturbation in context. To predict a perturbation in a test context, it takes as its prompt the training cells that received the same perturbation, in any training context, and as its query 500 control cells of the test context. It then generates the query cells as they would appear under that perturbation. It has no perturbation-identity input, so it applies only to perturbations that have training cells. We train it from scratch for each fold with the authors’ two-stage recipe, masked-gene pretraining followed by in-context post-training, using only the fold’s training and validation experimental conditions. We do not use the released weights because their gene vocabulary covers only 4,607 of the 6,642 Replogle–Nadig genes. As the authors do to stabilize training, we lower the peak pretraining learning rate to 3 × 10^−5^. We use 128 cells per cell set and select the checkpoint with the lowest validation loss.
- *Perturbation mean* (Adduri et al., 2026) predicts the control mean of the test context plus the perturbation’s pseudobulk delta expression averaged over the training contexts that measured it. It applies only to seen perturbations.
- *Context mean* (Adduri et al., 2026) predicts the control mean of the test context plus the mean pseudobulk delta expression of that context’s training perturbations, the same for every perturbation. For a context without training responses, it predicts the control mean.
- *Rhaister-O* (Svensson et al., 2026) is a zero-shot model. It predicts a perturbation’s DE statistics in a new context from that context’s control pseudobulk, through a per-perturbation, per-gene intercept and slope fit on the training DE statistics. Perturbations measured in too few training contexts fall back to the additive model of the same work. It applies only to seen perturbations.
- *COMPASS-N* (Liang and Singh, 2026) decomposes the training responses in the test context into shared axes, loadings, and residuals, and predicts an unseen perturbation from its nearest neighbours in the SPACE protein embedding (Hu et al., 2025), with 5 neighbours for the loadings and 10 for the residual. Perturbations without a SPACE embedding receive the training mean. It requires training responses in the test context and a gene target.
- *PRESAGE* (Littman et al., 2025) embeds each perturbation from 40 published gene knowledge graphs and two co-expression graphs computed from the training responses, and is trained per context. It requires training responses in the test context and a gene target.
- *TabICL* (Palla et al., 2026) uses the frozen TabICL regressor in two modes. For unseen perturbations, it regresses the principal components of pseudobulk delta expression on those of perturbation features, with the training responses of the test context as the in-context examples. The features are CellProfiler morphology, BioGPT, ESM2, STRING, and DepMap for genetic perturbations, and PIE’s small-molecule Knowledge Sources (perturbation text, SMILES, L1000, PRISM, and JUMP) for *Tahoe-100M*; a missing source enters as zeros. For unseen contexts, it maps perturbed cells of the training contexts to the test context by optimal-transport matching of control cells, which uses no perturbation features.

#### B.3. Generalization within dataset

In all three datasets, the validation set is 10% of each training context’s training experimental conditions, and PIE is trained at learning rate 10^−4^ for 5,000 optimizer steps. Test-set sizes below are for unseen contexts, unseen perturbations, and both unseen, in that order.

##### Replogle–Nadig (Replogle et al., 2022; Nadig et al., 2025)

The four cell lines contribute 611,211 cells. Each of four folds has one test cell line and 1,259–1,273 randomly drawn training perturbations, and every validation perturbation keeps a training response. The test sets hold 867–983, 1,172–1,252, and 386–470 experimental conditions and, with the training and validation sets, cover every experimental condition exactly once. There are 17 delta-expression bins (Appendix A.5), or 13 when RPE1 is the test line. The selected checkpoints lie at steps 1,518–2,631, and we report the mean and sample standard deviation across folds.

**Jiang et al**. **(2025)**. The 30 contexts are six cell lines (A549, BxPC-3, HAP1, HT-29, K562, MCF7) under five stimuli (IFN-*β*, IFN-*γ*, insulin, TGF-*β*, TNF-*α*). On-target knockdown filtering leaves 461 experimental conditions over 146 perturbations. Each of six folds has one test cell line, whose five contexts are the test contexts, and 92–102 randomly drawn training perturbations. The test sets hold 21–76, 92–128, and 6–43 experimental conditions. The selected checkpoints lie at steps 800–2,171. A fold’s score averages the contexts of its test set, and we report the mean and sample standard deviation across folds. Spearman effect size and Discrimination Score L1 are undefined in a context with one test perturbation.

##### *Tahoe-100M* (Zhang et al., 2025)

The 50 contexts are cancer cell lines, and the 1,137 perturbations are 379 small molecules at 0.05, 0.5, and 5 *µ*M, with DMSO-treated cells as controls. A single fold has five test cell lines and 265 training molecules at all three doses (795 perturbations), giving 32,173 training and 3,588 validation experimental conditions and test sets of 3,377, 13,055, and 1,451. The SMILES embedding and the measured compound profiles replace the gene-target Knowledge Sources (Appendix A.2). The selected checkpoint is at step 4,914. We report the mean and sample standard deviation across cell lines: the five test lines for unseen contexts and both unseen, and the 45 training lines for unseen perturbations.

##### Baselines

Context mean is evaluated in all three regimes and is the only baseline when both entities are unseen.

- *S**tate* is trained per fold on the cells of the training experimental conditions and the control cells of all contexts, for 50,000 steps at learning rate 10^−4^. We select the checkpoint with the lowest validation loss on Replogle–Nadig and use the final checkpoint on Jiang et al. It is evaluated on unseen contexts and not run on *Tahoe-100M*.
- *S**tack* is trained per fold on Replogle–Nadig, on all 6,642 genes, and evaluated on unseen contexts.
- *Perturbation mean* is computed from each fold’s training set and evaluated on unseen contexts.
- *Rhaister-O* is fit per fold on the training DE statistics, without a validation set, and evaluated on unseen contexts. Its full model, rather than the fallback, covers about a third of the Replogle–Nadig test experimental conditions, none of Jiang et al., and 5 of 3,377 of *Tahoe-100M*.
- *COMPASS-N* has no trainable parameters and is evaluated on unseen perturbations in Replogle–Nadig and Jiang et al.; it is not run on *Tahoe-100M*, whose perturbations have no gene target.
- *PRESAGE* is trained once per training context and fold, with early stopping on validation loss and, on Replogle–Nadig, the per-cell-line learning rates of the original paper. It is evaluated on unseen perturbations in Replogle–Nadig and Jiang et al. and, like COMPASS-N, not run on *Tahoe-100M*.
- *TabICL* is evaluated on unseen perturbations and unseen contexts in all three datasets. For the Jurkat context of Replogle–Nadig, only the STRING and DepMap features are available. On *Tahoe-100M*, it is fit once per training cell line on PIE’s small-molecule Knowledge Sources. Neither mode applies when both entities are unseen.

Tables 3–5 give the scores of all three datasets.

**Table 3.** Within-dataset generalization, unseen contexts. Mean ± sample standard deviation across folds (Replogle– Nadig, four; Jiang et al., six) or, for *Tahoe-100M*, across the five test cell lines of its single fold. Bold marks the best mean per dataset and metric; –, unable to perform the task; NaN, the metric is undefined. Higher is better for all metrics.

| Model | AUPRC | Jaccard | DE direction match | Spearman effect size | Spearman LFC | Discrimination Score L1 |
| --- | --- | --- | --- | --- | --- | --- |
| *Replogle–Nadig* |  |  |  |  |  |  |
| STATE | 0.138 ± 0.033 | 0.094 ± 0.024 | 0.622 ± 0.026 | 0.362 ± 0.385 | 0.244 ± 0.040 | 0.600 ± 0.017 |
| STACK | 0.175 ± 0.036 | 0.136 ± 0.034 | 0.680 ± 0.014 | 0.656 ± 0.109 | 0.358 ± 0.065 | 0.641 ± 0.071 |
| Perturbation mean | 0.088 ± 0.022 | 0.079 ± 0.018 | 0.806 ± 0.031 | 0.440 ± 0.106 | 0.534 ± 0.029 | 0.659 ± 0.073 |
| Context mean | 0.085 ± 0.023 | 0.076 ± 0.035 | 0.000 ± 0.000 | NaN | NaN | 0.502 ± 0.000 |
| Rhaister-O | 0.344 ± 0.028 | 0.173 ± 0.011 | 0.805 ± 0.032 | **0.741 ± 0.053** | 0.591 ± 0.060 | 0.712 ± 0.072 |
| COMPASS-N | – | – | – | – | – | – |
| PRESAGE | – | – | – | – | – | – |
| TabICL | 0.074 ± 0.019 | 0.075 ± 0.019 | 0.719 ± 0.029 | 0.633 ± 0.091 | 0.428 ± 0.055 | 0.631 ± 0.052 |
| PIE | **0.424 ± 0.034** | **0.185 ± 0.023** | **0.819 ± 0.026** | 0.734 ± 0.052 | **0.664 ± 0.046** | **0.764 ± 0.061** |
| *Jiang et al.* |  |  |  |  |  |  |
| STATE | 0.084 ± 0.038 | 0.082 ± 0.083 | 0.525 ± 0.079 | 0.218 ± 0.176 | 0.252 ± 0.160 | 0.588 ± 0.020 |
| Perturbation mean | 0.020 ± 0.010 | 0.018 ± 0.009 | 0.538 ± 0.032 | –0.032 ± 0.220 | 0.350 ± 0.152 | 0.586 ± 0.023 |
| Context mean | 0.022 ± 0.012 | **0.119 ± 0.102** | 0.000 ± 0.000 | NaN | NaN | 0.569 ± 0.026 |
| Rhaister-O | 0.186 ± 0.060 | 0.094 ± 0.066 | 0.578 ± 0.047 | 0.344 ± 0.179 | 0.280 ± 0.191 | 0.599 ± 0.022 |
| COMPASS-N | – | – | – | – | – | – |
| PRESAGE | – | – | – | – | – | – |
| TabICL | 0.017 ± 0.009 | 0.022 ± 0.011 | 0.443 ± 0.047 | –0.137 ± 0.228 | 0.199 ± 0.093 | 0.584 ± 0.011 |
| PIE | **0.214 ± 0.065** | 0.077 ± 0.042 | **0.704 ± 0.045** | **0.371 ± 0.153** | **0.523 ± 0.130** | **0.647 ± 0.039** |
| *Tahoe-100M* |  |  |  |  |  |  |
| Perturbation mean | 0.367 ± 0.136 | 0.264 ± 0.074 | 0.744 ± 0.021 | 0.307 ± 0.068 | 0.478 ± 0.058 | 0.813 ± 0.048 |
| Context mean | 0.373 ± 0.132 | 0.000 ± 0.000 | 0.000 ± 0.000 | NaN | NaN | 0.501 ± 0.000 |
| Rhaister-O | 0.665 ± 0.125 | 0.372 ± 0.045 | 0.684 ± 0.022 | **0.830 ± 0.131** | 0.403 ± 0.029 | 0.970 ± 0.019 |
| TabICL | 0.302 ± 0.122 | 0.375 ± 0.130 | 0.582 ± 0.028 | –0.049 ± 0.121 | 0.372 ± 0.038 | 0.866 ± 0.042 |
| PIE | **0.680 ± 0.094** | **0.400 ± 0.112** | **0.872 ± 0.053** | 0.759 ± 0.090 | **0.721 ± 0.104** | **0.982 ± 0.013** |

**Table 4.** Within-dataset generalization, unseen perturbations. Mean ± sample standard deviation across folds (Replogle–Nadig, four; Jiang et al., six) or, for *Tahoe-100M*, across the 45 training cell lines of its single fold. Bold marks the best mean per dataset and metric; –, unable to perform the task; NaN, the metric is undefined. Higher is better for all metrics.

| Model | AUPRC | Jaccard | DE direction match | Spearman effect size | Spearman LFC | Discrimination Score L1 |
| --- | --- | --- | --- | --- | --- | --- |
| *Replogle–Nadig* |  |  |  |  |  |  |
| STATE | – | – | – | – | – | – |
| STACK | – | – | – | – | – | – |
| Perturbation mean | – | – | – | – | – | – |
| Context mean | 0.082 ± 0.009 | 0.063 ± 0.011 | 0.675 ± 0.025 | NaN | 0.372 ± 0.035 | 0.503 ± 0.000 |
| Rhaister-O | – | – | – | – | – | – |
| COMPASS-N | 0.092 ± 0.010 | 0.083 ± 0.009 | 0.748 ± 0.009 | 0.336 ± 0.022 | 0.438 ± 0.024 | 0.645 ± 0.008 |
| PRESAGE | 0.091 ± 0.010 | 0.080 ± 0.009 | 0.745 ± 0.022 | 0.370 ± 0.071 | 0.444 ± 0.025 | 0.633 ± 0.014 |
| TabICL | 0.094 ± 0.010 | 0.085 ± 0.010 | 0.769 ± 0.022 | 0.442 ± 0.025 | 0.469 ± 0.036 | 0.682 ± 0.010 |
| PIE | **0.304 ± 0.023** | **0.168 ± 0.011** | **0.775 ± 0.022** | **0.525 ± 0.030** | **0.552 ± 0.022** | **0.709 ± 0.006** |
| *Jiang et al.* |  |  |  |  |  |  |
| STATE | – | – | – | – | – | – |
| Perturbation mean | – | – | – | – | – | – |
| Context mean | 0.026 ± 0.004 | 0.024 ± 0.004 | 0.632 ± 0.030 | 0.051 ± 0.110 | 0.273 ± 0.054 | 0.623 ± 0.010 |
| Rhaister-O | – | – | – | – | – | – |
| COMPASS-N | 0.026 ± 0.004 | 0.024 ± 0.004 | 0.613 ± 0.027 | 0.060 ± 0.106 | 0.263 ± 0.081 | 0.623 ± 0.011 |
| PRESAGE | 0.025 ± 0.004 | 0.024 ± 0.004 | 0.625 ± 0.036 | –0.032 ± 0.084 | 0.290 ± 0.059 | 0.624 ± 0.008 |
| TabICL | 0.026 ± 0.004 | 0.024 ± 0.004 | 0.601 ± 0.033 | –0.027 ± 0.072 | 0.195 ± 0.054 | 0.630 ± 0.009 |
| PIE | **0.112 ± 0.027** | **0.058 ± 0.019** | **0.707 ± 0.019** | **0.240 ± 0.133** | **0.501 ± 0.058** | **0.640 ± 0.013** |
| *Tahoe-100M* |  |  |  |  |  |  |
| Perturbation mean | – | – | – | – | – | – |
| Context mean | 0.306 ± 0.135 | 0.222 ± 0.093 | 0.630 ± 0.068 | –0.104 ± 0.166 | 0.287 ± 0.149 | 0.502 ± 0.000 |
| Rhaister-O | – | – | – | – | – | – |
| TabICL | 0.309 ± 0.136 | 0.221 ± 0.091 | 0.630 ± 0.071 | –0.041 ± 0.082 | 0.273 ± 0.128 | 0.519 ± 0.008 |
| PIE | **0.528 ± 0.109** | **0.316 ± 0.080** | **0.682 ± 0.094** | **0.031 ± 0.082** | **0.454 ± 0.159** | **0.531 ± 0.010** |

**Table 5.**
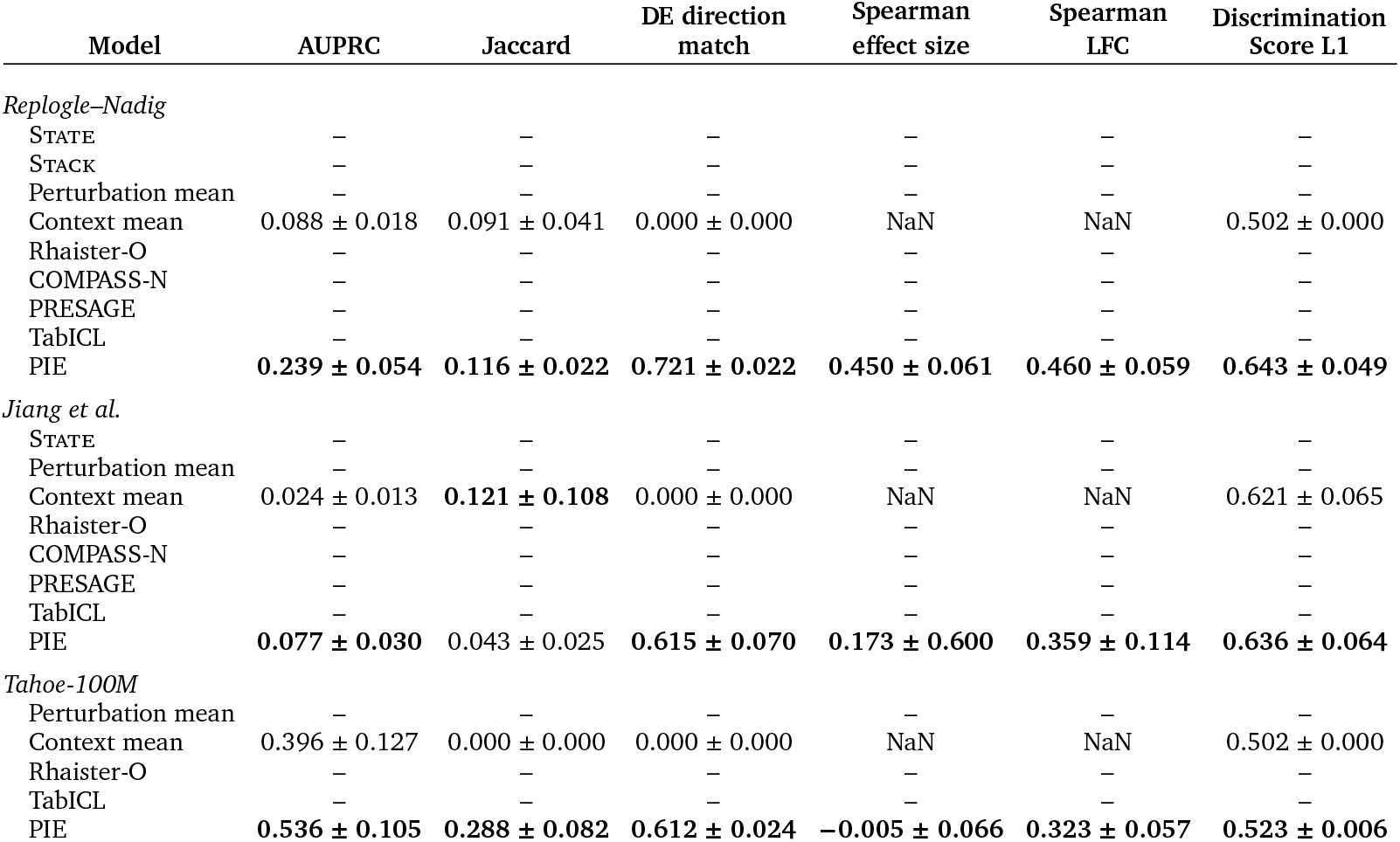
Within-dataset generalization, both unseen. Mean ± sample standard deviation across folds (Replogle–Nadig, four; Jiang et al., six) or, for *Tahoe-100M*, across the five test cell lines of its single fold. Bold marks the best mean per dataset and metric; –, unable to perform the task; NaN, the metric is undefined. Higher is better for all metrics.

#### B.4. Generalization across datasets

In every held-out-dataset experiment, PIE is trained on the other four datasets for 10,000 optimizer steps at learning rate 10^−4^ with a global batch of 128 experimental conditions and validated every 100 steps. The held-out dataset contributes no training or validation response, and contexts of a held-out cell line are removed from the training and validation sets. A context derived from a held-out cell line is discarded as well. Validation AUPRC averages over the contexts of each training dataset and then with equal weight over the four datasets. A perturbation is seen if it is measured in any context of any training dataset. The delta-expression bins have width ln 1.01, with the clipping range set by the largest 99.9th percentile among the training datasets (Appendix A.5). Test scores are computed per context of the held-out dataset, and we report the mean and sample standard deviation across contexts. Test-set sizes below are for seen and unseen perturbations, in that order.

##### Replogle–Nadig held out

Training samples *Tahoe-100M* (weight 0.68; 51,144 experimental conditions, 45 cell lines, 379 small molecules, 1,137 perturbations), X-Atlas/Orion (0.30; 12,628, two lines, 9,679 perturbations), Jiang et al. (0.01; 342, 23 contexts, 139 perturbations), and Arc VCC 2025 (0.01; 146, one context). Removed: the five Jiang et al. K562 contexts and *Tahoe-100M* Hep-G2/C3A, derived from HepG2. Validation: 957 experimental conditions (*Tahoe-100M* 224 from four lines, X-Atlas/Orion 656, Jiang et al. 33 from two HT-29 contexts, Arc VCC 2025 44). Test: 2,286 (723 perturbations) and 3,173 (1,134), with 1,253, 1,439, 1,314, and 1,453 in HepG2, Jurkat, K562, and RPE1. Bins: 121; selected step: 4,700.

##### Jiang et al. held out

Training samples *Tahoe-100M* (0.584), X-Atlas/Orion (0.258), Replogle–Nadig (0.15), and Arc VCC 2025 (0.009): 65,597 experimental conditions, 49 contexts, 11,952 perturbations. Removed: Replogle–Nadig K562 and *Tahoe-100M* A-549 and HT-29. Validation: 1,172. Test: 227 (71 perturbations, 27 contexts) and 234 (75, 29). Bins: 155; selected step: 9,700.

##### Atlas/Orion held out

Training samples *Tahoe-100M* (0.826), Replogle–Nadig (0.15), Jiang et al. (0.012), and Arc VCC 2025 (0.012): 56,921 experimental conditions, 78 contexts, 3,161 perturbations; nothing removed. Validation: 613. Test: 841 (775 perturbations) and 12,443 (9,232), with 8,987 in HCT116 and 4,297 in HEK293T. Bins: 147; selected step: 8,300.

##### Arc VCC 2025 held out

Training samples *Tahoe-100M* (0.584), X-Atlas/Orion (0.258), Replogle–Nadig (0.15), and Jiang et al. (0.009): 69,403 experimental conditions, 79 contexts, 12,017 perturbations; nothing removed. Validation: 1,225. Test: 121 and 69 in the single H1 context, so no standard deviation is reported. Bins: 147; selected step: 8,600.

##### Baselines

In every experiment, perturbation mean, context mean, Rhaister-O, and TabICL are evaluated on seen perturbations, and context mean alone on unseen perturbations.

- *Perturbation mean* averages, for each gene, the perturbation’s pseudobulk delta expression over the training contexts, from any training dataset, that measure it and the gene.
- *Context mean* predicts the control mean of the held-out context, since no training response is measured there.
- *Rhaister-O* is fit on the DE statistics of the training set. Its full model covers no test experimental condition in any experiment, so every prediction comes from the fallback.
- *TabICL* uses its unseen-context mode, with the training contexts that measure the test context’s seen perturbations as sources.

Tables 6 and 7 give the scores for all four held-out datasets.

**Table 6.** Generalization across datasets, seen perturbations. Mean ± sample standard deviation across the contexts of each held-out dataset: four cell lines for Replogle–Nadig, two for X-Atlas/Orion, and 27 cell line–stimulus contexts for Jiang et al.; Arc VCC 2025 has a single context and no standard deviation. Bold marks the best mean per dataset and metric; –, unable to perform the task; NaN, the metric is undefined. Higher is better for all metrics.

| Model | AUPRC | Jaccard | DE direction match | Spearman effect size | Spearman LFC | Discrimination Score L1 |
| --- | --- | --- | --- | --- | --- | --- |
| *Replogle–Nadig* |  |  |  |  |  |  |
| Perturbation mean | 0.062 ± 0.013 | 0.052 ± 0.010 | **0.733 ± 0.034** | 0.439 ± 0.077 | 0.472 ± 0.041 | 0.561 ± 0.026 |
| Context mean | 0.064 ± 0.014 | 0.086 ± 0.043 | 0.000 ± 0.000 | NaN | NaN | 0.502 ± 0.000 |
| Rhaister-O | 0.283 ± 0.048 | 0.130 ± 0.029 | 0.730 ± 0.034 | 0.547 ± 0.101 | 0.476 ± 0.048 | 0.582 ± 0.034 |
| TabICL | 0.052 ± 0.011 | 0.051 ± 0.011 | 0.634 ± 0.030 | 0.452 ± 0.087 | 0.272 ± 0.084 | 0.540 ± 0.016 |
| PIE | **0.378 ± 0.033** | **0.138 ± 0.009** | 0.727 ± 0.035 | **0.627 ± 0.064** | **0.508 ± 0.043** | **0.620 ± 0.055** |
| *Jiang et al.* |  |  |  |  |  |  |
| Perturbation mean | 0.025 ± 0.023 | 0.019 ± 0.018 | 0.497 ± 0.099 | −0.017 ± 0.478 | 0.187 ± 0.167 | 0.581 ± 0.058 |
| Context mean | 0.027 ± 0.025 | **0.123 ± 0.172** | 0.000 ± 0.000 | NaN | NaN | 0.579 ± 0.045 |
| Rhaister-O | 0.073 ± 0.048 | 0.086 ± 0.119 | 0.445 ± 0.081 | **0.197 ± 0.502** | 0.141 ± 0.174 | 0.585 ± 0.051 |
| TabICL | 0.023 ± 0.022 | 0.026 ± 0.025 | 0.431 ± 0.175 | −0.071 ± 0.556 | 0.139 ± 0.276 | 0.582 ± 0.070 |
| PIE | **0.162 ± 0.069** | 0.055 ± 0.078 | **0.615 ± 0.115** | 0.180 ± 0.494 | **0.425 ± 0.269** | **0.586 ± 0.050** |
| *X-Atlas/Orion* |  |  |  |  |  |  |
| Perturbation mean | 0.016 ± 0.008 | 0.015 ± 0.008 | 0.620 ± 0.054 | 0.190 ± 0.111 | 0.405 ± 0.064 | 0.560 ± 0.024 |
| Context mean | 0.012 ± 0.007 | 0.134 ± 0.089 | 0.000 ± 0.000 | NaN | NaN | 0.504 ± 0.004 |
| Rhaister-O | 0.207 ± 0.046 | **0.145 ± 0.032** | 0.650 ± 0.055 | 0.576 ± 0.070 | 0.471 ± 0.081 | 0.571 ± 0.032 |
| TabICL | 0.010 ± 0.006 | 0.010 ± 0.006 | 0.590 ± 0.023 | 0.135 ± 0.206 | 0.376 ± 0.004 | 0.568 ± 0.039 |
| PIE | **0.307 ± 0.070** | 0.080 ± 0.000 | **0.809 ± 0.073** | **0.628 ± 0.086** | **0.709 ± 0.065** | **0.657 ± 0.045** |
| *Arc VCC 2025* |  |  |  |  |  |  |
| Perturbation mean | 0.281 | 0.124 | 0.549 | 0.076 | 0.132 | 0.524 |
| Context mean | 0.289 | 0.000 | 0.000 | NaN | NaN | 0.506 |
| Rhaister-O | 0.299 | 0.005 | 0.550 | 0.536 | 0.133 | 0.524 |
| TabICL | 0.271 | **0.202** | **0.571** | 0.429 | 0.142 | 0.509 |
| PIE | **0.395** | 0.076 | 0.559 | **0.604** | **0.147** | **0.678** |

**Table 7.** Generalization across datasets, unseen perturbations. Mean ± sample standard deviation across the contexts of each held-out dataset: four cell lines for Replogle–Nadig, two for X-Atlas/Orion, and 29 cell line–stimulus contexts for Jiang et al.; Arc VCC 2025 has a single context and no standard deviation. Bold marks the best mean per dataset and metric; –, unable to perform the task; NaN, the metric is undefined. Higher is better for all metrics.

| Model | AUPRC | Jaccard | DE direction match | Spearman effect size | Spearman LFC | Discrimination Score L1 |
| --- | --- | --- | --- | --- | --- | --- |
| *Replogle–Nadig* |  |  |  |  |  |  |
| Perturbation mean | – | – | – | – | – | – |
| Context mean | 0.102 ± 0.026 | 0.077 ± 0.034 | 0.000 ± 0.000 | NaN | NaN | 0.502 ± 0.000 |
| Rhaister-O | – | – | – | – | – | – |
| TabICL | – | – | – | – | – | – |
| PIE | **0.252 ± 0.061** | **0.123 ± 0.018** | **0.603 ± 0.034** | **0.560 ± 0.029** | **0.278 ± 0.089** | **0.515 ± 0.009** |
| *Jiang et al.* |  |  |  |  |  |  |
| Perturbation mean | – | – | – | – | – | – |
| Context mean | 0.024 ± 0.037 | **0.110 ± 0.170** | 0.000 ± 0.000 | NaN | NaN | **0.578 ± 0.050** |
| Rhaister-O | – | – | – | – | – | – |
| TabICL | – | – | – | – | – | – |
| PIE | **0.089 ± 0.077** | 0.019 ± 0.017 | **0.629 ± 0.136** | **0.066 ± 0.425** | **0.341 ± 0.256** | 0.577 ± 0.075 |
| *X-Atlas/Orion* |  |  |  |  |  |  |
| Perturbation mean | – | – | – | – | – | – |
| Context mean | 0.001 ± 0.001 | **0.409 ± 0.082** | 0.000 ± 0.000 | NaN | NaN | 0.500 ± 0.000 |
| Rhaister-O | – | – | – | – | – | – |
| TabICL | – | – | – | – | – | – |
| PIE | **0.065 ± 0.016** | 0.173 ± 0.066 | **0.510 ± 0.141** | **0.096 ± 0.055** | **0.502 ± 0.064** | **0.504 ± 0.002** |
| *Arc VCC 2025* |  |  |  |  |  |  |
| Perturbation mean | – | – | – | – | – | – |
| Context mean | 0.196 | 0.000 | 0.000 | NaN | NaN | 0.510 |
| Rhaister-O | – | – | – | – | – | – |
| TabICL | – | – | – | – | – | – |
| PIE | **0.273** | **0.031** | **0.541** | **0.409** | **0.088** | **0.516** |

#### B.5. Ablating signal components

##### Shapley values

Let *N* be the set of *n* = 5 players. A subset *S* ⊆ *N* is a set of players present in a model, with the other players removed, and *v* (*S*) is the test metric of the model trained with exactly the players in *S*. The Shapley value of player *i* sums over the subsets that exclude *i* (Shapley, 1953):

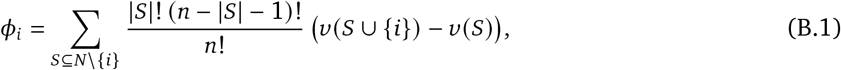

and the values satisfy the efficiency property of the same work,

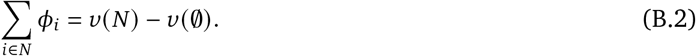

In the experiment of Section 4.4, *v* (*∅*) is the model with all five groups removed. We compute *ϕ*_*i*_ exactly in each fold, where the values add up to that fold’s difference.

##### Ablation of Perturbation Knowledge Sources

A second experiment attributes the metrics to the five Perturbation Knowledge Sources individually: the Perturbation Description, NCBI Gene Annotations, ESM2, STRING, and DepMap. The Context Description and the three evidence channels are present in every model, so the model without any source is the first experiment’s model with only the Perturbation Knowledge Sources removed. The protocol is otherwise that of Section 4.4.

##### Removal

A removed input is absent in training and at test. A removed Knowledge Source contributes no tokens, and a sample for which no source remains receives the zero carrier token (Section 3.2). Removing Control Mean Evidence drops the control channel from the per-gene evidence, and removing Same-context or Same-pert Evidence sets that channel and its flag to zero in every training and test sample, the encoding of a channel with no contributing response (Section 3.3).

##### Models and training

Each experiment has 2^5^ = 32 models per fold, and the two experiments share two: the full model, and the model without Perturbation Knowledge Sources, which is the first experiment’s model with only that group removed. This gives 62 models per fold and 248 in total. Each model follows the recipe of Section 4.2 with one seed and at most 5,000 optimizer steps, and the checkpoint with the highest validation AUPRC is used for all metrics and regimes. Every model stops early once validation AUPRC has not improved by 0.001 for 1,000 optimizer steps; the median model stops at step 3,282 (range 1,822 to 5,000, with 22 models reaching the step limit).

### Results

The five sources add the most AUPRC for unseen perturbations, raising it from 0.281 to 0.306 (Figure 7). Of this gain, DepMap contributes 0.013, STRING 0.008, and NCBI Gene Annotations 0.006; ESM2 contributes 0.000 and the Perturbation Description −0.002. For Discrimination Score L1 in this regime, every source is positive, from 0.020 for ESM2 to 0.066 for DepMap, and the five together account for 0.206. For unseen contexts, the sources add 0.013 AUPRC, most of it from ESM2 (0.007) and DepMap (0.004). With both entities unseen, the sources leave AUPRC unchanged (−0.001). They raise DE direction match by 0.075, with every source contributing between 0.010 and 0.021, Spearman LFC by 0.101, and Discrimination Score L1 by 0.143. The Perturbation Description has a negative AUPRC value in all three regimes.

**Figure 7.**
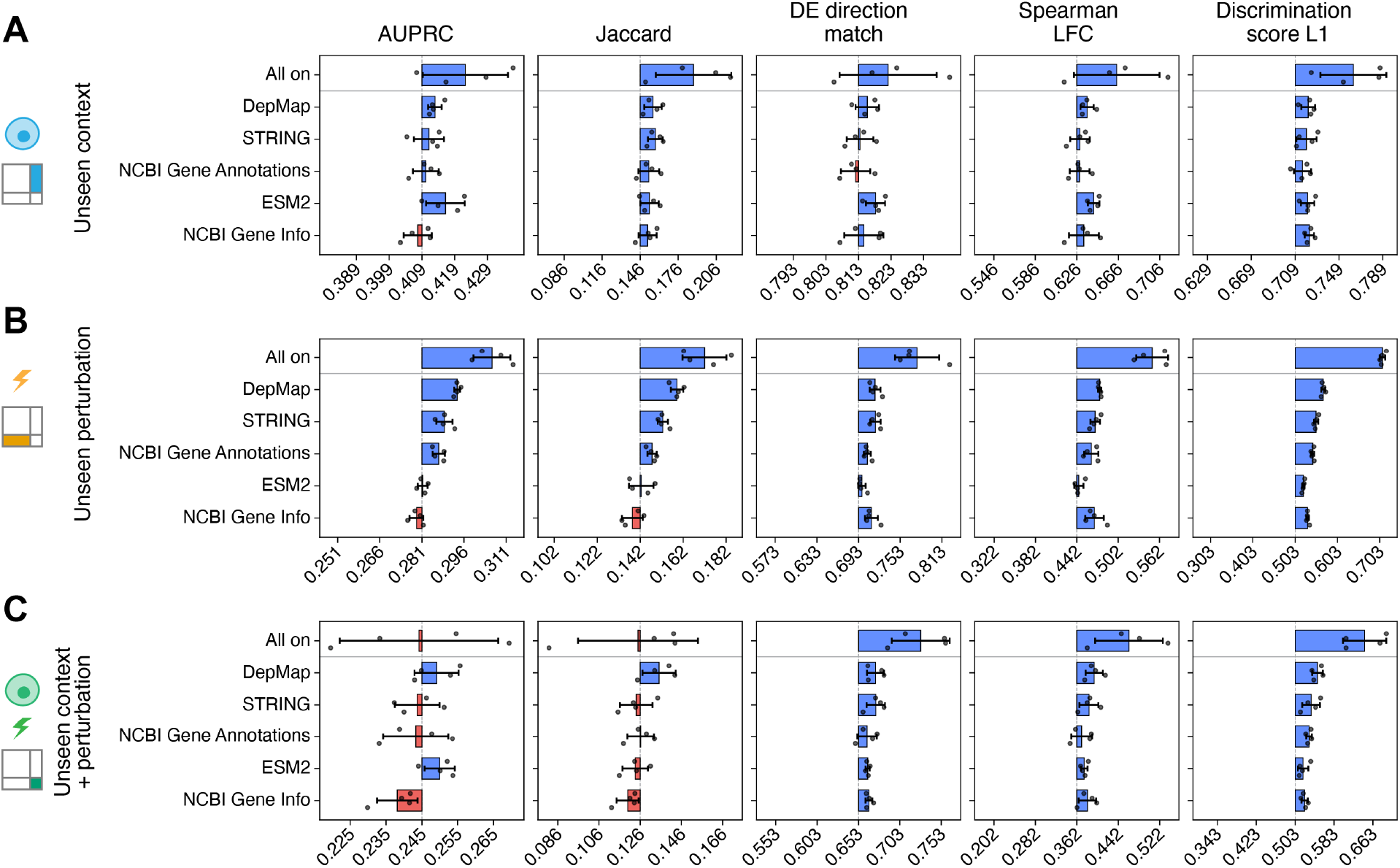
Contribution of Perturbation Knowledge Sources. Shapley values of the five Perturbation Knowledge Sources on the within-dataset task, computed from models trained with every subset of the sources: the Perturbation Description (NCBI Gene Info), NCBI Gene Annotations, ESM2, STRING, and DepMap, with the other four input groups present in every model. Rows are the three regimes. Each axis is centred (dashed line) on the model without any source, and the top bar (All on) is the difference between the models with all and with no sources; tick labels are metric values. Bars show means and std across four folds, each holding out one cell line; dots are folds. DepMap contributes the most AUPRC among the sources for unseen perturbations.

